# HDX-MS performed on BtuB in E. coli outer membranes delineates the luminal domain’s allostery and unfolding upon B12 and TonB binding

**DOI:** 10.1101/2022.01.07.475440

**Authors:** Adam M. Zmyslowski, Michael C. Baxa, Isabelle A. Gagnon, Tobin R. Sosnick

## Abstract

To import large metabolites across the outer membrane of Gram-negative bacteria, TonB dependent transporters (TBDTs) undergo significant conformational change. After substrate binding in BtuB, the *E. coli* vitamin B12 TBDT, TonB binds and couples BtuB to the inner membrane proton motive force that powers transport ^(1)^. But, the role of TonB in rearranging the plug domain to form a putative pore remains enigmatic. Some studies focus on force-mediated unfolding ^(2)^ while others propose force-independent pore formation ^(3)^ by TonB binding leading to breakage of a salt bridge termed the “*Ionic Lock*”. Our hydrogen exchange/mass spectrometry measurements in *E. coli* outer membranes find that the region surrounding the Ionic Lock, far from the B12 site, is fully destabilized upon substrate binding. A comparison of the exchange between the B12 bound and the B12&TonB bound complexes indicates that B12 binding is sufficient to unfold the Ionic Lock region with the subsequent binding of a TonB fragment having much weaker effects. TonB binding accelerates exchange in the third substrate binding loop, but pore formation does not obviously occur in this or any region. This study provides a detailed structural and energetic description of the early stages of B12 passage that provides support both for and against current models of the transport process.

**Significance Statement:** TonB dependent transporters such as BtuB are found in the outer membranes of Gram-negative bacteria. They import scarce nutrients essential for growth, such as B12, the substrate of BtuB. Many transport steps remain enigmatic. Recent studies have emphasized force-mediated unfolding or the breakage of the “Ionic Lock”, a moiety far from the B12 binding site. A strong dependence on the membrane environment has been noted. Accordingly, we measured hydrogen exchange on BtuB still embedded in native outer membranes and found that B12 binding is sufficient to break the Ionic Lock. The amino terminus then extends into the periplasm to bind TonB. But we find no evidence of pore formation, which likely requires energy transduction from the inner membrane by TonB.

## Introduction

TonB-dependent transporters (TBDTs) are ubiquitously found in the outer membranes (OMs) of Gram-negative bacteria where they import scarce nutrients ^(4)^. Some members of the family function as virulence factors, counteracting nutrient sequestration by the innate immune response ^(5)^. Aspects of the transport mechanism remain unclear, especially the nature of the structural changes involved in the creation of the pore required for substrate passage through the transmembrane β-barrel ^(6)^. Pore formation in each TBDT is tailored to its substrate, which can range in size from 56 Da (iron) to the 1.4 kDa cyanocobalamin (B12). In some TBDTs, the large extracellular loops close behind the substrate. This closure prevents back-diffusion ^(7)^ implying that formation of a pore through the lumen of the barrel need not imply bi-directional diffusion. Alternatively, the pore may form in stages, with no single conformation allowing for unrestricted diffusion across the length of the membrane. Though the individual mechanisms may be substrate-specific, pore formation generally involves conformational changes within the plug domain that normally occludes the lumen.

BtuB is a prototypical *E. coli* TBDT that has been studied extensively in both functional and biophysical contexts **(Figure 1)** ^(1, 8, 9)^. B12 is a relatively large TBDT substrate, which places constraints on the minimum pore size. In proposed models of pore formation ^(2, 3)^, B12 binding initiates an allosteric signaling event that leads to the release of the 7-residue, amino terminal “Ton box” ^(10)^ into the periplasm where it binds the periplasmic carboxy terminal domain (CTD) of TonB (TonB_CTD_), an inner membrane (IM) protein ^(11)^.

**Figure 1.**
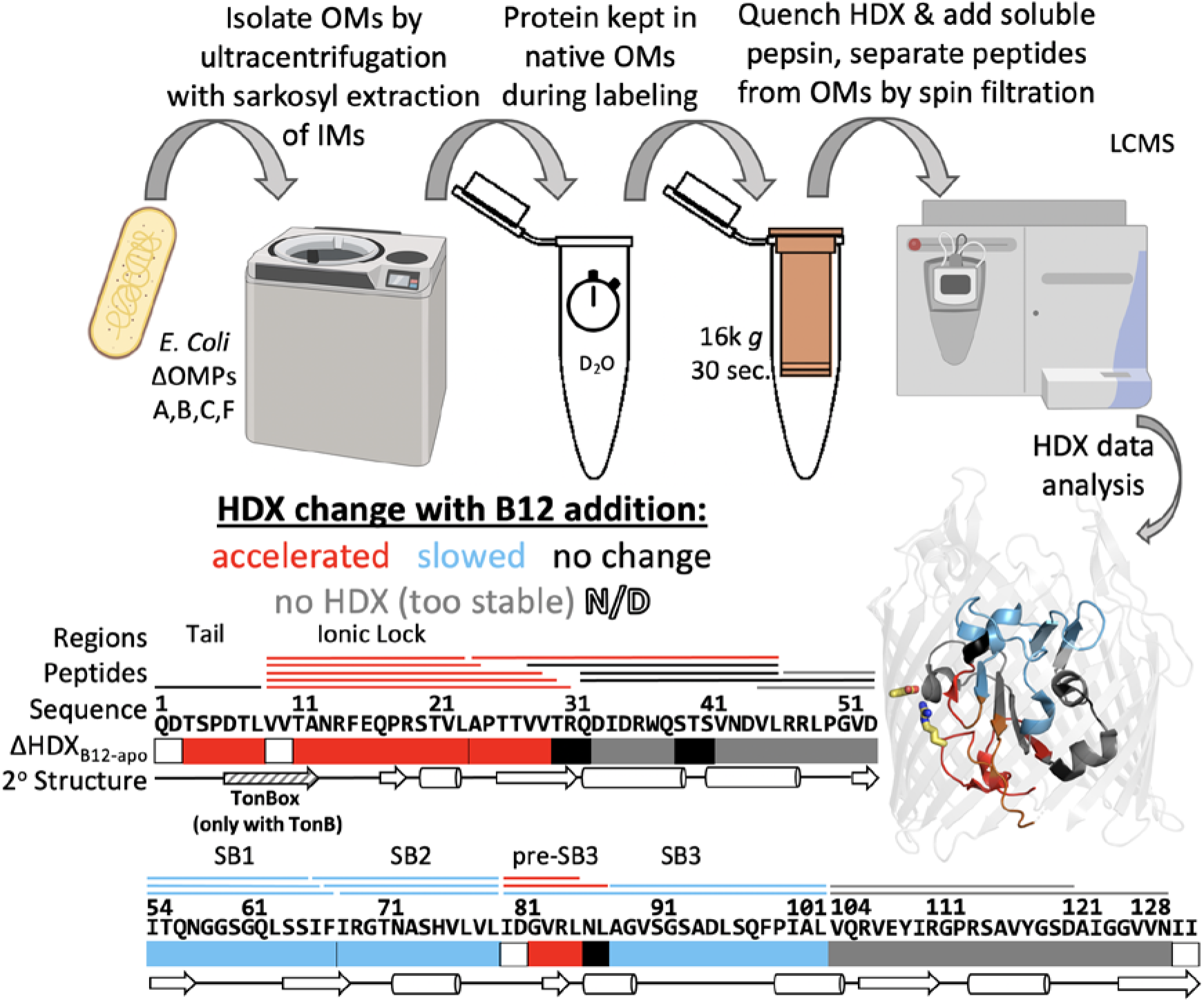
BtuB purification and HDX protocol. Sample preparation in outer membranes (OMs) involves no chromatography, only repeated centrifugation and resuspension with a sarkosyl extraction step to remove inner membranes (IMs). All steps except HDX labeling occur on ice. Structure shown is for the apo state of BtuB (PDBID: 1NQE), with the β-barrel domain displayed transparently to highlight regions of the plug domain.

The steps leading from TonB_CTD_-Ton box binding to the formation of a pore are currently under debate. The binding of TonB_CTD_ enables the energetic, and possibly mechanical, coupling between the IM and OM that is necessary for transport against a B12 gradient ^(12)^. In the periplasm, TonB interacts with the ExbB/ExbD complex, which harvests the proton motive force of the IM ^(13)^. Though many structures of the TonB/ExbB/ExbD complex have recently been solved, the relationship between the transport mechanism and the assumed rotation of this inner membrane complex on TBDTs remains ambiguous^(6)^. In force-dependent models, TonB transduces energy from the IM either by pulling the plug domain’s amino terminus towards the periplasm ^(2, 8, 14)^ or by applying a torque ^(15)^. These forces are proposed to remodel the plug to form a pore, as seen in simulations ^(2, 16)^. Other steered MD simulations have characterized the interactions of B12 as it was pulled through the lumen ^(17)^. Alternatively, pore formation has been proposed to be regulated in a force-independent manner by the *Ionic Lock (IL)*, a conserved salt bridge between Arg14 on the plug and Asp316 on the nearby inner surface of the transmembrane β-barrel ^(3, 18)^. In the force-independent model, TonB binding need only disengage the *IL* to allow pore-forming motions in the third substrate binding loop (SB3) when B12 is present ^(3)^.

In the B12-bound crystal structure of BtuB (PDBID 1NQH ^(9)^), the *IL* region adopts a very similar backbone conformation to the apo state (1NQE). The major change in the *IL* region is the rotation of the Arg14 sidechain and the loss of the salt bridge with Asp316. **(Figure S1)** In the ternary complex with TonB_CTD_, the *IL* is fully broken and the Arg14 C_α_ atom moves by 30.2 Å (2GSK, ^(8)^). This breakage occurs along with the residues immediately carboxy terminal to the Ton box adopting an extended conformation, which permits them to exit the lumen and bind TonB_CTD_ **(Figure S1)**.

Cafiso and coworkers have suggested that the folded *IL* region in B12-bound BtuB (1NQH) is a feature resulting from crystallographic solutes and packing contacts perturbing the folding equilibrium ^(19)^. This potentially crucial difference leaves unresolved the actual conformation of the *IL* region in the B12 bound state in the native membrane. Determining when the *IL* region unfolds is important as the subsequent binding of TonB has recently been proposed to be the critical event that breaks the *IL* rather than the binding of B12 ^(3)^. This ambiguity on how the amino terminal region of BtuB responds to B12 and subsequent TonB binding events is the focus of the present investigation.

Studying the forces that promote transport in the context of the intact cell envelope is desirable but so far has not been accomplished. Hence, most biophysical measurements of BtuB have been performed in reconstituted, single bilayer systems. However, as recently emphasized ^(3)^, the results of these experiments have depended on the choice of environment, which have included detergents, liposomes, supported bilayers, OMs, or recently even live *E. coli* ^(20)^. The pore-forming intermediate state of BtuB has proven difficult to characterize, and efforts have relied on either extraction from native membranes, the use of cysteine mutants and spin labels, or a combination thereof ^(2, 20)^.

Here we present a label-free investigation of the effects of B12 and TonB_CTD_ binding on the conformation of BtuB plug domain using hydrogen-deuterium exchange (HDX) in OMs **(HDX-MS, Figure 1)**. As anticipated, we find that binding of B12 slows down HDX on the plug’s three substrate binding (SB) loops **(Figure 1, blue)**. Upon the binding of B12 alone, however, HDX is *increased* by 10^2^-10^3^-fold (i.e., destabilized) for at least a dozen residues surrounding the *IL*. These residues stretch from the carboxy terminal side of the Ton box to the beginning of the first β-strand **(Figure 1, red)**, supporting the proposal that a binding event breaks the *IL* as a prerequisite to transport ^(3)^. However, we find that the binding of B12 alone, rather than TonB, disrupts the *IL*. Accordingly, we propose that B12 binding leads to an allosterically-induced unfolding of BtuB’s amino terminus via breakage of the *IL*, which in turn enables TonB to bind the newly-exposed Ton box **(Figure 1, red)**. In addition, we observe a mild acceleration within a four-residue segment located between the *IL* and the B12-contacting apex of the SB3 loop, which is near a site of pore formation recently proposed by Cafiso and coworkers ^(3)^. This co-localization suggests that B12 initiates only the first step in a cascade of events that promotes its own transport **(Figure 2)**. The binding of TonB is a subsequent step, but our results suggest that it too may be insufficient to cause pore formation in the absence of coupling to an active ExbB/ExbD complex. We end with a discussion of the similarity to prior findings ^(3)^ but also note some differences.

**Figure 2.**
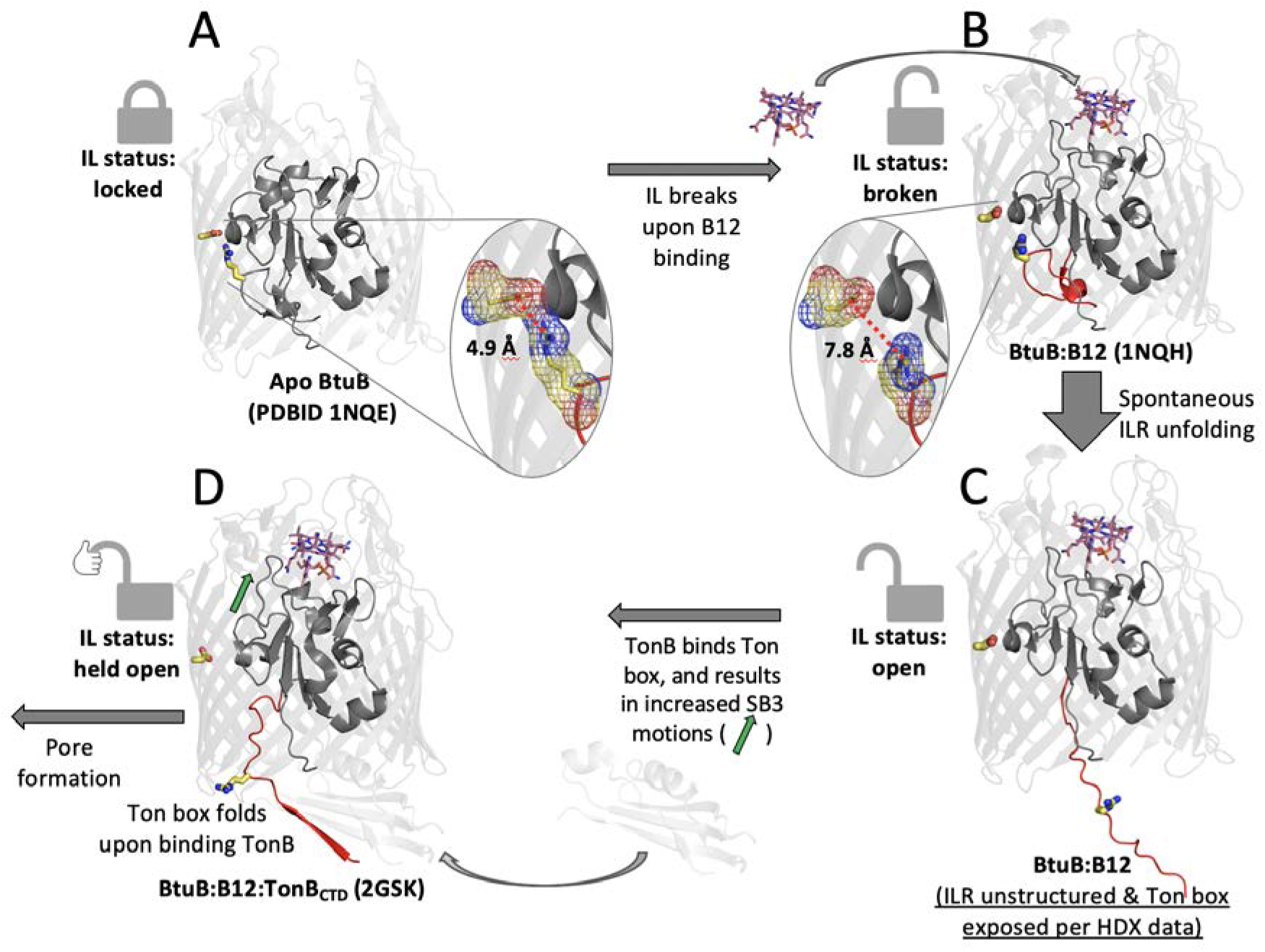
Model of the BtuB plug domain’s conformational response to two ligand binding events. **A, C, D)** are the three major steps identified by the HDX data whereas **B)** is a structure with bound B12 observed by crystallography but not observed in our data. The *Ionic Lock* sidechains are shown as yellow sticks, with oxygen and nitrogen atoms colored in red and blue, respectively. Inset ovals show an expanded mesh representation of the *IL* in states A) and B).

## Results

We first discuss the protocols and reproducibility of our HDX measurements in OMs, followed by a presentation of the data for regions where stability increases due to B12 binding, focusing on the substrate binding loops SB1-SB3 **(Figure 3, shades of blue/green)**. We next describe the large decrease in stability observed near the *IL* and the smaller stability losses for regions at the nearby amino terminus, the first strand, and the start of the SB3 loop **(Figure 3, red shading)**. We conclude with an examination of the regions where minimal changes in HDX are seen upon B12 binding. To allow access to the HDX data of this study, we are including HDX data summary **(Table 1)**, HDX peptide summary **(Table S1)**, and HDX data tables **(Dataset 1)** ^(21)^.

**Table 1.** Biochemical and statistical details for each measured state.

| Data Set | apo BtuB | BtuB + B12 | BtuB + TonB | BtuB + B12 + TonB (Ternary) |
| --- | --- | --- | --- | --- |
| HDX reaction details | 46.7 mM Na<sub>2</sub>HPO<sub>4</sub>, 0.6 mM Tris, pD<sub>read</sub> = 6.80, 22 °C | 46.7 mM Na<sub>2</sub>HPO<sub>4</sub>, 0.6 mM Tris, pD<sub>read</sub> = 6.80, 20 µM B12, 1 mM CaCl<sub>2</sub>, 22 °C | 46.7 mM Na<sub>2</sub>HPO<sub>4</sub>, 0.6 mM Tris, pD<sub>read</sub> = 6.80, 22 °C | 46.7 mM Na<sub>2</sub>HPO<sub>4</sub>, 0.6 mM Tris, pD<sub>read</sub> = 6.80, 2 µM B12, 1 mM CaCl<sub>2</sub>, 22 °C |
| HDX time course (min) | 0.1, 0.316, 1.0, 3.16, 10, 31.6, 100, 316, 1620 | 0.1, 0.316, 1.0, 3.16, 10, 31.6, 100, 316, 1620 | 0.1, 0.33, 1.0, 3.5, 10, 36 | 0.1, 0.316, 1.0, 3.16, 10, 31.6, 100, 316 |
| HDX control samples | Maximally-labeled control (LH4 construct, his-tagged on extracellular loop) | Maximally-labeled control (LH4 construct, his-tagged on extracellular loop) | Maximally-labeled control (LH4 construct, his-tagged on extracellular loop) | Maximally-labeled control (LH4 construct, his-tagged on extracellular loop) |
| Back-exchange (mean / IQR) | 25.1% / 9.0% | 25.1% / 9.0% | 25.1% / 9.6% | 24.9% / 9.3% |
| # of Peptides | 35 | 35 | 29 | 31 |
| Sequence coverage | 95.4% (of 130 residues covering N-terminus & plug domain)<br>21.2% (whole protein) | 95.4% (of 130 residues covering N-terminus & plug domain)<br>21.2% (whole protein) | 90% (of 130 residues covering N-terminus & plug domain)<br>19.7% (whole protein) | 90% (of 130 residues covering N-terminus & plug domain)<br>19.7% (whole protein) |
| Average peptide length / Redundancy | 16.5 / 0.7 | 16.5 / 0.7 | 16.5 / 0.7 | 15.9 / 0.6 |
| Replicates | 3 (biological) | 3 (biological) | 1 | 3 (biological) |
| Repeatability | 2% / 0.24 Da (average standard deviation) | 2% / 0.24 Da (average standard deviation) | N/A | 3.2% / 0.37 Da (average standard deviation) |
| Significant differences in HDX (ΔHDX > X Da) | 1.14 Da (98% confidence interval) | 1.16 Da (98% confidence interval) | N/A | 2.66 Da (98% confidence interval) |

**Figure 3.**
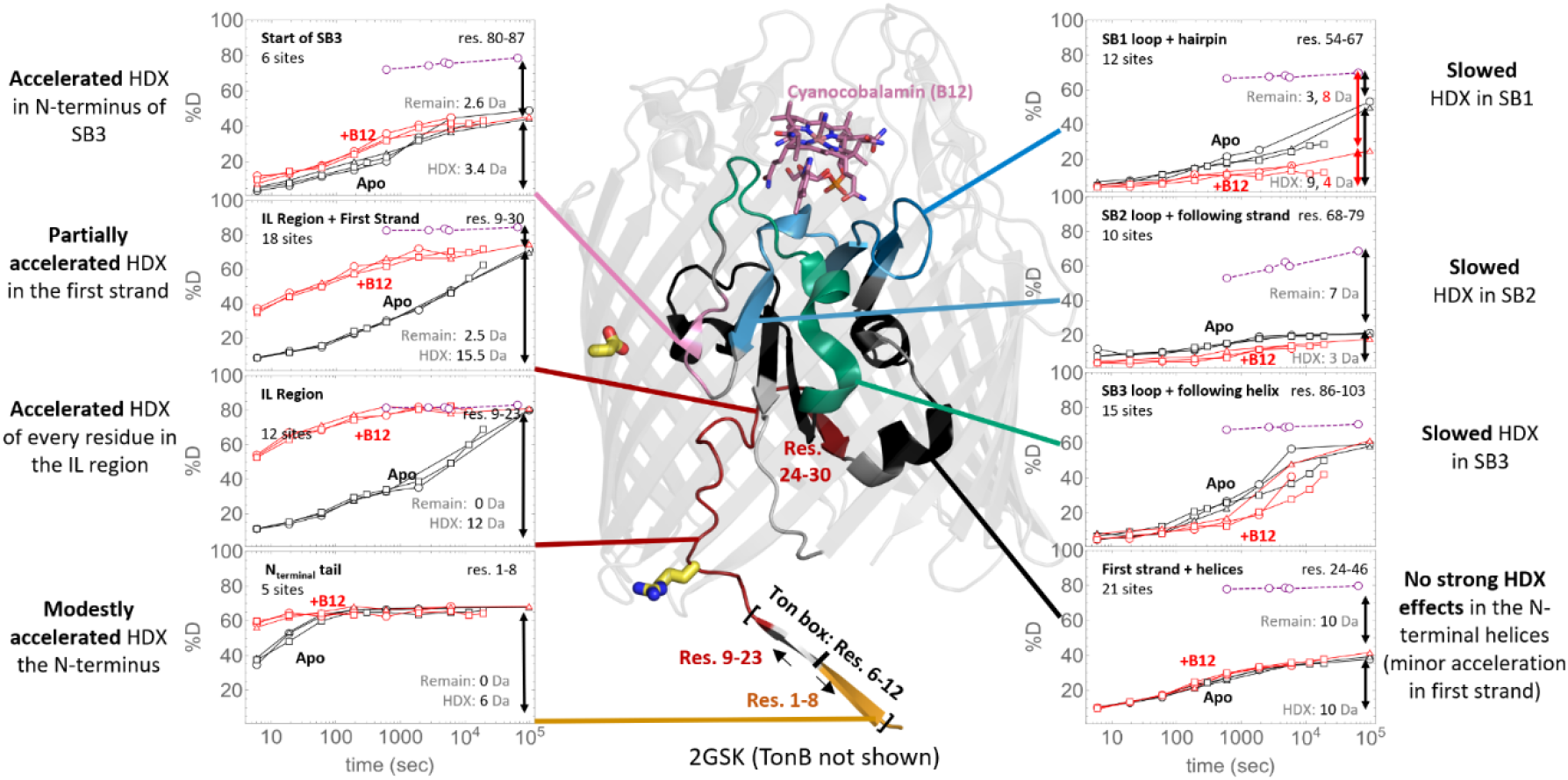
Effects of B12 binding on HDX. Uptake plots show biologically triplicated data (circles, triangles, and squares) for BtuB measured in apo (black) and B12-bound (red) states. Insets contain uptake curves for the peptides depicted by the colored regions on the BtuB structure. The number of exchangeable sites is assumed to be the number of non-proline residues minus two. The exchanged and remaining masses at the last time point are given in grey, after correcting for back exchange by normalizing to non-native controls which assumed to be fully deuterated (purple circles with dashed lines).

### OM sample preparation, sequence coverage

We measured deuterium uptake at pD 7.2 (pD_read_ = 6.8), 22 °C for apo, B12-and TonB_CTD_-liganded states of BtuB in labeled in native-like OMs from three biological replicates. To study the effects of TonB_CTD_ binding, we used TonB_ΔN_, a construct containing the CTD and periplasmic linker, but lacking the amino terminal transmembrane helix (hereafter referred to as TonB within the context of our experiments). OMs and OM proteins (OMPs) are significantly more stable than their IM counterparts, and thus the protocols used to study OMPs employ strategies that are not as commonly used for IM proteins.

To our knowledge, the only published HDX-MS study of OMPs performed in a native OM-like environment was conducted on OmpF in *E. coli* OM vesicles ^(22)^. BtuB, however, did not express well in OM vesicles, so we adapted protocols used in the EPR studies of BtuB embedded in native-like OMs ^(23)^. Based on a purification strategy employed on FhuA ^(24)^, a BtuB construct was created with five histidines, one glycine, and two serines inserted after His449 on the apex BtuB’s seventh extracellular loop. This construct and the wild-type protein were overexpressed in

*E. coli*, and after lysis, total membranes were separated using ultracentrifugation followed by extraction with sarkosyl to selectively remove the IMs. Initially, the OMPs were extracted with detergent, but detergent-solubilized BtuB failed to exhibit evidence of B12 binding in the *IL* region by HDX, consistent with earlier findings that detergent solubilization perturbs the folding of the amino terminus ^(25)^. Hence, BtuB was left in its native OM environment for our HDX studies.

Accordingly, our workflow was altered to allow HDX labeling as well as digestion of BtuB still embedded in the OM preparations. After HDX labeling, we added denaturant and soluble pepsin in the low pH quench buffer rather than employing an in-line protease column as typically done in HD-XMS measurements. This change in protocol afforded us greater control over the cleavage conditions while eliminating fouling of the chromatographic system by OM components such as lipids. We optimized digestion for redundant and nearly complete peptide coverage of the plug domain, in part by decreasing β-barrel cleavage **(Figures 1, S2B)**. Consistent with the known trends of BtuB unfolding in response to denaturant, ^(26)^ the proteolysis was best near the amino-terminus. Pepsin’s weak preference to cleave after hydrophobic residues caused the peptides to cluster into regions with similar boundaries with peptides within each region exhibiting similar HDX behavior. We use a nomenclature where regions are capitalized while the sequence bounds are denoted in subscript, with important peptides in parentheses, e.g., BtuB’s amino terminus is covered by Region_3-8_ (Peptide_1-8_).

The imperfect correspondence between peptide and region boundaries is caused by an effect common to all HDX-MS studies. The intrinsic exchange rates for an exposed amide (*k*_chem_) for the two amino-terminal residues are sufficiently fast that the they largely lose their deuterium labels via back-exchange, and hence, they do not measurably impact the deuterium uptake level ^(27)^. Consequently, we defined regions to start at the third residue of the most amino terminal peptide of the cluster except for situations where other peptides can provide information for those residues. Our partitioning of the plug domain into regions typically coincided with the secondary structure boundaries. Areas previously reported to undergo distinct motions usually separated into distinct regions **(Figure 3)** ^(28)^. Despite each region having markedly different exchange behavior, peptides within a region were internally consistent with respect to their deuteration levels. Peptides spanning more than one region displayed intermediate effects, weighted by the number of amide protons in each region **(Figure S6)**. While we have obtained peptide coverage sufficient to calculate site-resolved uptake at some positions, this is not generally possible. Accordingly, the analysis was conducted at the peptide level across the multiple replicates and conditions.

### Reproducibility and heterogeneity

Three biological replicates were initially prepared. One replicate used unmodified BL12 (DE3) *E. coli* whereas two replicates used a quadruple OMP knockout strain to examine the potential effects of endogenous OMPs ^(29)^. The use of the knockout strain did not significantly alter the HDX results **(Figures 3-4)**, although it did reduce the non-BtuB OMP content as desired **(Figure S2A)**. One of the two knockout replicates had high levels of non-native signals, complicating the analysis. Although the slowly exchanging subpopulations matched the other samples, this replicate was excluded from analysis to simplify data processing and a fourth knockout sample was prepared to take its place.

For most peptides, the slowest exchanging population was dominant and hence, was considered to represent the natively folded protein. For certain peptides, however, we observed multiple populations with distinct exchange kinetics **(Figure S3)**. Their isotopic envelopes remained separated even in our longest measurements of 10^5^ seconds, implying that the underlying populations do not interconvert on this time scale. The heterogeneity generally was manifested as bimodal isotopic envelopes that varied in relative intensity for different peptides but exhibited a systematic correlation within a bioreplicate.

Given the observed heterogeneity, we utilized the bimodal analysis capability of HDExaminer 3.1 program despite the additional complexity. We note that the use of a unimodal analysis does not qualitatively change the major findings **(Figure S3)** but it does introduce systematic differences between bioreplicates. These, however, are resolved with bimodal analysis, which provides additional motivation for taking advantage of the software’s bimodal capability.

When minor populations were present, they usually exchanged much faster than the major population, implying that they are less structured or stable. Several additional features such as their insensitivity to B12 suggested that these fast populations represent non-native or damaged proteins **(Figures S3-S4, S7)**. The uptake trends of the minor populations were similar to membraneless controls measured in 0.5 M urea. These controls were prepared using a BtuB construct that lacked a signal sequence and were expressed in inclusion bodies and purified in 8 M urea so that the β-barrel was unlikely to have ever been properly folded. It should be noted, however, that the control samples still exhibited substantial protection relative to *k*_chem_ (see SI HDX formalism section), and thus we did not feel justified in using this control for estimating *k*_chem_ **(Figure S7**, see discussion). By comparing our main dataset to this control, we identified and excluded populations of peptides that likely arose from a subset of non-native BtuB molecules and focused our analysis on the major, slowly exchanging population(s). The minor subpopulations often had overlapping mass spectral envelopes for at least some time points, and so were not always computationally isolated or suppressed using the bimodal fitting capability of the HDExaminer 3.1 program. These features were not plotted in the main figures **(see Figures S3-S4** and **Table S1)** but are recorded in **Dataset 1**. We additionally required consistency between multiple peptides and replicates in constructing our comprehensive profile of the plug domain’s behavior.

Our conclusions regarding B12 binding hold for every presented peptide across all replicates, though minor features of HDX uptake trends varied between bio-replicates **(Figure 4)**. For the major population, the only significant difference between biological replicates is that one showed a diminished effect of TonB_ΔN_ binding **(Figure 4B)**. Optimization of integration bounds for our data resulted in the apo state’s deuterium level having a standard deviation of ∼0.24 Da when averaged across the 35 presented peptides. **Dataset 1** provides estimates of mass differences significant at the 98% confidence level for each peptide. These intervals varied from 0.17 (Peptide_45-53_) to 8.30 Da (Petide_80-103_) and had a mean of 1.14 Da and median of 0.62 Da. The average level of back exchange observed in our data was 25%, with an interquartile range of 9.0%. Further plots summarizing the reproducibility are provided in **Figure S5**.

**Figure 4.**
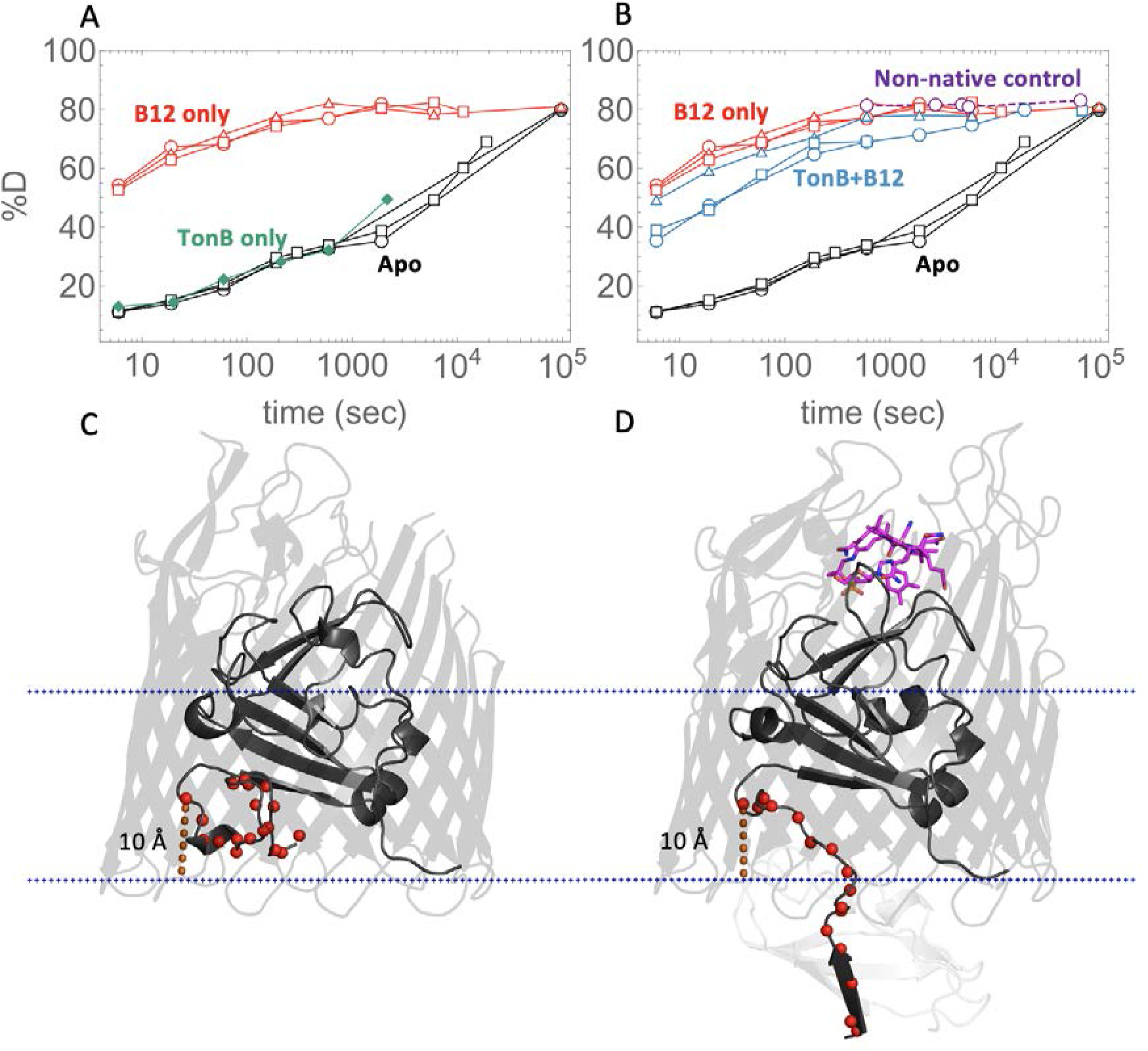
HDX in the *Ionic Lock* Region. **A, B)** Uptake plots for Peptide_9-23_, three biological replicates (black), and (red), and TonB_ΔN_ (green). Four long points of an non-native control shown in purple. B) Uptake plots for Peptide_9-23_, apo (black), B12 (red), and B12+TonB_ΔN_ (blue) **C) & D):** Position of the amino terminal regions relative to the membrane plane. **C)** apo (1NQE) and **D)** BtuB:B12:TonB complex (2GSK) crystal structures, taken from the OPM database ^(32)^. N atoms for residues 6-23 are shown as red spheres. Distances from Leu23’s N atom to the nearest membrane plane dummy atom are shown as dashed orange lines, in both cases about 10 Å.

### Substrate binding loops

In apo-BtuB, substrate binding loops SB1-SB3 exchanged slowly with most associated peptides retaining greater than 50% of their H-level even after 10^5^ seconds, the longest measured time point. This slow exchange translates to an HDX protection factor PF > 10^5^, where PF is defined as the relative slowing of the observed HDX rate (*k*_obs_) compared to *k*_chem_: PF *= k*_chem_*/k*_obs_ (see SI for HDX formalism and inference of EX2 behavior). Exchange was further slowed in the presence of 20 μM B12. An examination of overlapping peptides indicated that this slowing was concentrated in the residues nearer to the B12 binding site. Sites in SB1 had the largest decrease in fractional deuterium uptake, followed by SB2 and SB3 **(Figure S6)**. In the presence of B12, the exchange rates for the few observable sites on SB2 were about 10-fold slower, although most sites already exchanged very slowly in the apo state making any changes difficult to discern **(Figure S6.15-S6.24)**. Whereas the overall exchange for SB3 was slowed only 5-fold, some peptides displayed complex behavior, and exchange for other peptides was even accelerated, as will be discussed later. Otherwise, the general slowing of HDX upon the addition of B12 in the three SB loops indicates that B12 binding stabilizes regions directly contacting the substrate.

### Allosteric destabilization of the *Ionic Lock* and Ton box upon B12 binding

Our major finding is that upon B12 binding, exchange drastically increased for the *IL* Region_11-23_ (Peptides_9-23, 9-27, 9-28, 9-29, 9-30_). The *IL* region contains the Arg14-Asp316 salt bridge that connects the plug domain to the inner wall of the β-barrel **(Figures 4, S6.2-S6.6)**. In addition, exchange was mildly increased for the amino terminal tail Region_3-8_ **(Figure S6.1)**. Peptides in the *IL* region exchanged up to 10^3^-fold faster with B12 with a majority of the exchange occurring within the 6 second dead-time of the measurement. For the tail region, deuterium uptake increased mildly, with the uptake increasing from 50% to 80% after 6 seconds. From this widespread acceleration of HDX, we infer that both regions are unfolded in the B12 bound state and that upon B12 binding the amino terminus partially exits the lumen. The smaller change in HDX for the tail region is the result of it being less stable in the apo state to begin with.

We interpret the sizable B12 effect on the HDX of the *IL* region to result from the breaking of interactions between residues within this region and the β-barrel, most importantly the Arg14-Asp316 salt bridge. This disruption results in the Ton box’s exit into the periplasm. Residues 1, 2, 9, and 10 are the first two residues on each of their respective peptides and thus unobservable due to back exchange (an inherent HDX-MS issue occurring during sample handling). These four residues likely are also unstructured given the rapid exchange of every observable residue surrounding them (e.g., Peptides_1-8 and 9-23_). Notwithstanding, the data provide strong evidence that BtuB’s first 23 residues undergo a complete or near-complete loss of structure in response to B12 binding. One might suppose the opposite situation where a sizable population of folded *IL* remains after B12 addition but exchanges via localized fluctuations for residues not encompassing the entire *IL* region; however, this scenario should, upon the addition of TonB, shift the equilibrium to an increased unfolded *IL* population and a corresponding acceleration of HDX for at least part of the *IL* region. The opposite, however, was observed suggesting that the *IL* was broken for a majority of BtuB molecules in the presence of B12 alone.

Our inference that both the *IL* region and the Ton box were unfolded upon B12 binding alone is further supported by our finding that the addition of TonB_ΔN_ did not further accelerate exchange. The binding of TonB (specifically the carboxy terminal domain) to the Ton box requires that the Ton box be extended into the periplasm, which in turn requires that the *IL* be broken. Therefore, if the *IL* region remained folded after B12 binding and then unfolded when TonB_ΔN_ bound, one would have expected a marked increase in exchange for the *IL* region upon TonB_ΔN_ binding. However, exchange did not markedly increase upon the addition of TonB_ΔN_ (in fact, exchange even slowed for a few residues, which provides direct evidence of TonB binding) **(Figure S6.2-S6.6)**. This minimal increase in exchange is in the opposite direction of what would be expected if B12 binding did not cause the *IL* to break. Therefore, the *IL* region is already unfolded prior to TonB_ΔN_ binding, and B12 binding alone is sufficient to unfold the *IL* region.

In ostensible disagreement with this conclusion is that the *IL* region retains a significant amount of HDX protection even after B12 binding. The deuterium buildup curve is ∼10^2^ slower than what would be expected for exposed amides in bulk solution ^(27)^. To rationalize the observed slowing, we propose that the bulk solution rate may be an inappropriate reference in the present situation as the *IL* region’s carboxy terminus is located 10 Å inside the barrel lumen, which prevents portions of the region from fully exiting into bulk solvent. In this partially confined state inside the barrel, the access of OD^-^ to the peptide backbone also could be reduced and exchange slowed. The residues that do exit into the periplasm may still be subject to a milder but similar blocking effect from interactions with the nearby peptidoglycan **(Figure S7)**. In addition, the effective [OD^-^] should be lower in the barrel than in the bulk solvent as the anion prefers to be hydrated rather than reside in the lumen’s lower dielectric environment ^(30)^. Both the reduced access to the backbone and lower [OD^-^] concentration would retard exchange and may account for the slower than expected exchange observed upon B12 binding for the *IL* and for other regions, especially those further inside the barrel.

In the B12-bound state, Cafiso et al. proposed that the *IL* is broken and were able to infer residues 1-15 as being unstructured ^(10)^. Early EPR measurements probing the Ton box found that B12 binding caused the first residue within the Ton box, Asp6, to project 20-30 Å into the periplasm, leading the authors to model the first 15 residues as exiting the lumen with a broken *IL* ^(10)^. Consistently our HDX-MS measurements also identified this sizable unstructured region, finding that it extends out to residue 23. We observed rapid deuteration of every observable amide out to residue 23 (Peptides_1-8 and 9-23_) suggesting that an additional eight residues are affected by the same B12 binding event. Hence, we have evidence of a B12-initiated unfolding that is larger than previously described.

A comparison of Peptide_9-23_ with overlapping Peptides_9-27, 9-28, 9-29, and 9-30_ indicates that acceleration of HDX was also occurring in some additional residues between 24-29, which encompass BtuB’s first β-strand **(Figures S5.7-S5.12)**. This acceleration did not equally affect all five exchangeable residues, however. The exchange for some residues within BtuB’s first β-strand remained very slow, implying that the strand remains folded with its outer edge becoming more solvent exposed when B12 binds. In sum, we have used HDX to quantify the size of the B12-mediated unfolding in BtuB’s amino-terminus.

### Complex ligand effects in the third substrate binding loop, including destabilization by B12

In sharp contrast to the generally slowed exchange observed for the substrate binding loops, the amino terminus of SB3 (Region_82-85_) exhibited accelerated exchange upon B12 binding **(Figure S6.24)**. The apparent timescale of exchange for the fastest amide proton among residues 82-85 on Peptide_80-85_ was approximately 2,000 s, with B12 binding resulting in faster exchange rate of this single amide by ∼5-fold. Other amides within Region_82-85_ remained too protected to be observed exchanging in our experiments, so the data are agnostic on whether these positions were affected by B12, unlike the remaining portion of SB3 (Region_86-103_) and the other SBs, which were affected **(Figures S6.15-S6.23)**. The change in HDX for Region_82-85_ upon B12 binding was relatively mild compared to that seen at the amino terminus of the protein. This region, however, was unique in having its exchange further enhanced beyond the already B12-accelerated rates by the addition of TonB_ΔN_. (**Figure S6.24)**. Hence, the SB3 region allosterically responds to distal binding of B12 as well as TonB binding events, supporting its relevance to the TonB-dependent transport mechanism ^(3)^. Two reasons may explain the relatively small changes in HDX observed in the SB3 region. Possibly, one end of the SB3 loop is accelerated while the other end is slowed by binding B12, yielding a small net change in uptake. Additionally, rather than pore formation, the small changes may reflect a modest change in H-bonding across the region.

### Regions with no significant HDX differences

The HDX of the three remaining regions of the plug domain did not display a significant response to B12 binding **(Figures S5611-S6.13, S6.33-S6.35)**. Two of these regions (Regions_30-46 and 47-53_) lie between the *IL* and SB1 while the third region (Region_104-128_) is carboxy terminal to SB3. A comparison of overlapping peptides within Region_30-46_ **(Figures S6.11-S6.12)** confirmed the mild response to B12 within residues 26-29, as discussed earlier, demonstrated that the residues flanking BtuB’s first stable helix do not respond to B12. Regions_47-53 and 104-128_ **(Figures S6.33-S6.35)** exhibited almost no exchange, being extremely stable. The high overall stability for these regions and apparent lack of a response to ligand binding suggest that rearrangements in the plug domain during ligand binding are restricted to the areas surrounding the *IL* and SB3. Strictly speaking, however, we must be agnostic on potential changes for residues that exchange outside our experimental time window.

## Discussion

Our major observation is that the binding of B12 promotes the complete breaking of the *Ionic Lock* ^(30)^, a critical salt bridge connecting the amino terminal end of the plug domain to the inner surface of the β-barrel (Arg14-Asp316). The breakage results in all of the residues within Region_11-23_ unfolding, exposing the Ton box and permitting TonB binding. Upon subsequent TonB binding to the B12-bound state, we did not observe evidence of pore formation or further unfolding of the *IL* region. In fact, TonB binding alone only produced subtle HDX changes throughout the plug domain, suggesting that its full function in pore formation requires an energized inner membrane, or some other factor only present in live cells.

Binding of B12 alone resulted in stability changes throughout the plug domain, both increasing and decreasing HDX rates. We observed the anticipated slowing of exchange for the loops that directly contact B12. But the addition of B12 also resulted in a dramatic acceleration of HDX by up to 10^3^-fold for all amide protons in the *IL* Region (e.g., Peptide_9-23_), even approaching the rapid exchange rates seen in BtuB’s labile amino terminus (Peptide_1-8_). The large increase in HDX rates upon B12 binding for residues in the Ton box reflects the disruption of the entire *IL* region (11-23) and can occur in response to B12 binding alone. We believe this disruption is responsible for exposing the Ton box, thereby regulating TonB binding.

The *IL* is over 20 Å away from the B12 binding site, implying an allosteric mechanism is involved in the breaking of this salt bridge. The amino terminal segment of SB3, Region_82-85_, is located near the *IL* and is also accelerated by B12 binding, pointing to the involvement of SB3 in the allosteric mechanism. Additionally, HDX of Region_82-85_ is further accelerated by the binding of TonB to the Ton box, suggesting a bi-directional coupling of one or both Ton box-containing regions with the SB3 loop. The movement of SB3 upon B12 binding likely is a critical initial step in the signal propagation downwards to the *IL*. Overall, our data support Cafiso’s proposal that SB3 may undergo further motions upon TonB binding ^(3)^.

As discussed earlier, even with a 10^2^-10^3^-fold increase in exchange rate for the *IL* Region_11-23_ upon binding B12, HDX exchange is still 10^2^-fold slower than the intrinsic exchange rate *k*_chem_. One might consider that this residual protection is indicative of some apparent structure in the *IL* region. However, evidence exists that this region is indeed unfolded: Because the binding of TonB_ΔN_ resulted in no further unfolding of the *IL* region after B12 binding, the *IL* must already have been broken and the region unfolded in the B12-bound state. Therefore, we propose that the 10^2^-fold residual protection arises from other factors including restricted access to the peptide backbone as the region is tethered inside the lumen as well as a reduction of the local [OD^-^] in the lumen’s lower dielectric environment ^(30)^. Additionally, H-bonding interactions with the peptidoglycan network, which normally undergirds the inner surface of the OM, may be conferring protection even to an otherwise unstructured chain. Further studies are needed to examine which, if any of the 3 mechanisms, is operational.

### Comparisons to existing models

Current models of BtuB transport can be broadly separated into two categories based on the proposed location and energetics of pore formation, including whether it is force-independent (FI) or force-dependent (FD). In both models, either implicitly or explicitly, the binding of B12 is necessary for exposure of the Ton box and subsequent TonB binding. In the FI model ^(3)^, TonB binding provides the energy for subsequent plug remodeling events, which are sufficient to open a pore near the *IL* region and permit B12 transport. By contrast, a recent FD model proposes that TonB binding establishes a mechanical linkage that transmits a force to unfold the “mechanically weak subdomain” within the plug ^(2)^. This partial unfolding of the plug forms a substrate channel, although some uncertainty exists regarding whether this mechanism applies *in vivo* ^(3)^.

While the results of our experiments do not fundamentally rule out either model, our HDX data demonstrate that binding of B12 alone is sufficient to disrupt the *IL* region and expose the Ton box. However, our data find that neither the individual B12 nor TonB binding events are sufficient to destabilize a substantial fraction of the plug domain, which is postulated to unfold under the FD model (Regions_31-46, 47-53, and 54-67_). Peptides in Region_45-53_ did not exchange in the duration of our measurements (10^5^ sec), implying a very high stability (Peptide_45-53_; PF > 10^7^). Flanking regions that unfold in the FD model (as observed in the atomic force microscopy study ^(2)^) were either unaffected (Peptide_31-46_; PF ∼10^4^) or even stabilized (Peptide_54-67_; PF ∼10^3^-10^6^) upon B12 binding. We believe that this high level of stability is unlikely, but not impossible, within the FD model ^(2)^. Our data also corroborate the argument ^(3)^ deployed against FD models related to the application of a torque on BtuB by TonB ^(15)^. We found that in the ternary complex of BtuB, B12, and TonB, nearly a dozen residues between the Ton box and BtuB’s stably folded core are unstructured, consistent with findings for FhuA ^(31)^. For these residues, the facile rotation of their (*ϕ,Ψ*) backbone torsion angles likely precludes the transmission of a torque directly from the Ton box to the plug.

Nevertheless, our HDX results also differ with the FI model regarding the effects of ligand binding. We find that the *IL* is broken by B12 binding alone, whereas Cafiso et al. ^(3)^ found *in vivo* that the *IL* is partially broken by B12 and only completely broken by TonB, based on the conformational response of the SB3 loop to B12 binding. Specifically, mutations that broke the *IL* (Arg14Ala, Asp316Ala) resulted in a 20 Å movement in SB3 and were reported to mimic TonB binding. In OMs, however, we find that the *IL* fully breaks in response to B12 binding, and additionally, that the *IL* is not broken by TonB binding at a concentration of 26 μM when B12 is absent. When B12 is added, however, we find the *IL* is sufficiently destabilized that subsequent TonB binding does not cause a significant slowing or acceleration of the HDX for the *IL* region. In our view, the Ton box is normally sequestered in the lumen of the β-barrel and becomes available for binding only when B12 binding allosterically induces the breakage of the *IL*. In this mechanism, the rest of the steps required to gate the pore cannot begin before the initial B12 binding event.

Whereas Cafiso et al.^(3)^ observed a 20 Å conformational change in SB3, we detected subtle HDX changes for this region upon B12 and TonB binding **(Figures S6.24 & S6.25)**. It is difficult to reconcile large scale motions occurring in SB3 with the lack of a large HDX effect for peptides covering the apex of SB3, especially since a 10^3^-fold increase was seen in the *IL* region. Possible explanations for the difference include our use of sarkosyl to remove IMs ^(3)^ and the EPR study’s use of live cells which differ from our OM preparation. Furthermore, the two techniques measure very different quantities and timescales from which protein motions are inferred. HDX is sensitive to H-bonding whereas EPR provides distance information from the spectral changes based on energy transfer between spin probes.

In our HDX-MS measurements, the introduction of TonB and formation of the ternary complex reduced the signal quality for certain peptides due to chromatographic overlap and mass spectral crowding. Interestingly, for certain SB3 peptides, deuterium uptake trends became complex even with B12 alone, i.e., tri-modal isotope envelopes **(Figure S4)**, possibly reflecting increased heterogeneity where a single population present in the apo state splits into multiple slowly interconverting populations upon ligand binding. Further study of the complex HDX behavior of the SB3 loops may shed additional light on the allosteric mechanism.

## Conclusions

Using HDX-MS, we identified regions of the BtuB plug domain that both increase and decrease stability upon substrate binding. Most notably, the binding of B12 alone is sufficient to initiate an allosteric pathway involving the disruption of a dozen residues located up to 20 Å away from the B12 binding site. The process involves the unfolding of the *IL* region to permit the binding of TonB to the Ton box.

The remaining steps in the transport process are less clear. Potentially, an exit channel forms that follows a route from the B12 binding site, past SB3, and then out through the space formerly occupied by the *IL* region. Alternatively, TonB binding may initiate the force-induced unfolding of the plug core to create a substrate channel. This latter mechanism is appealing in its simplicity and generalizability, as one can envision the role of substrate binding is simply to release the Ton box into solution where forces are applied by TonB, and the channel is then formed by the unfolding of approximately half of the plug domain core. However, the HDX data indicate that the core of the plug remains extremely stable when B12 and TonB are bound, providing a powerful example of how HDX’s ability to access local thermodynamic information can help identify mechanism. Besides the study of mutations that constitutively break the *IL* or disrupt potential allosteric pathways that cross the membrane, the measurement of the denaturant dependence of the HDX ^(27)^ could further define the conformational changes resulting from binding and force.

Because results across multiple techniques have depended on the membrane environment, future BtuB studies should be performed in as native-like a context as possible ^(3)^. Our application of HDX/MS in native OMs is a compromise between working in live cells, where transport intermediates may not accumulate due to the action of TonB, and in reconstituted systems such as detergents (which do not bind B12) or liposomes (which are compositionally very different from OMs). The use of OMs provides both a native-like bilayer and high yield, which may enable the ambitious goal of performing HDX *in vivo*.

## Materials and Methods

Outer membranes used for HDX experiments were obtained by French press lysis of *E. coli* overexpressing BtuB followed by several rounds of centrifugation and a brief extraction with sarkosyl to selectively remove inner membranes. HDX labeling was initiated by diluting concentrated suspensions of the outer membranes into deuterated solutions. HDX labeling was terminated and digestion simultaneously initiated by diluting the HDX labeling reaction into ice cold quench buffer (pH 2.5) previously supplemented with urea, soluble pepsin, microporous zirconium oxide beads, and the detergent DDM. After a brief incubation period, the labeled peptides released into solution were separated from insoluble undigested protein and membranes by spin filtration immediately before injection onto the HDX-LCMS system. Further details are described in SI Materials and Methods.

Chemicals were purchased from Sigma-Aldrich (St Louis, MO) unless otherwise noted in SI Materials and Methods. All BtuB and TonB constructs were expressed and purified as described in SI Materials and Methods.

## Acknowledgments

We thank R. Nakamoto, H. Hong, and N. Noinaj, E. Perozo, M. Clark, and members of the Sosnick lab for valuable conversations about TBDT biology, comments on the manuscript, or for kindly supplying plasmids and cell strains. This work was supported by NIH Research Grant R01 GM055694.

