## Supplementary material for "HDX-MS performed on BtuB in E. coli outer membranes delineates the luminal domain’s allostery and unfolding upon B12 and TonB binding": SI Appendix

### **Supplementary Information for**
HDX-MS maps the allosteric response and unfolding of BtuB's luminal domain to B12 and TonB binding in *E. coli* outer membranes

Adam M. Zmyslowski[1], Michael C. Baxa[1], Isabelle A. Gagnon[1], Tobin R. Sosnick[1-3]

Tobin R. Sosnick


**This PDF file includes:**

Supplementary text
Figures S1 to S7
Table S1
SI References

**Other supplementary materials for this manuscript include the following:**

Dataset S1: HDX data table (separate .xlsx file). HDX data and all computed quantities used the analysis.

**Supplementary text**

**HDX formalism.**

In apo-BtuB, substrate binding loops SB1-SB3 exchanged slowly with most associated peptides retaining greater than 50% of their H-level even after $10^5$ seconds, the longest measured time point. This slow exchange translates to an HDX protection factor PF > $10^5$, where PF is defined as the relative slowing of the observed HDX rate ($k_{obs}$) compared to the intrinsic chemical exchange rate for an exposed amide proton ($k_{chem}$): PF=$k_{chem}/k_{obs}$. In the commonly used formalism, hydrogen exchange is assumed to occur when an H-bond is broken in a transient "open state", and the NH is exposed to solvent [(1)],

$$\text{H bond formed} \underset{k_{close}}{\overset{k_{open}}{\rightleftarrows}} \text{H bond broken} \xrightarrow{k_{chem}} \text{H} \quad \text{D exchange}$$

where $k_{open}$ and $k_{close}$ are the opening and closing rates, respectively. The observed HDX rate for this reaction is given by

$$k_{obs} = \frac{k_{chem} \cdot k_{open}}{k_{close} + k_{open} + k_{chem}}.$$

Under the so-called EX2 condition where $k_{close} >> k_{chem}$, the PF is directly related to the equilibrium stability according to [(2)]:

$$\Delta G_{HX} = RT \ln K_{eq} = RT \ln\left(\frac{[closed]}{[open]}\right) = RT \ln\left(\frac{k_{close}}{k_{open}}\right) = RT \ln(PF - 1)$$

In the EX1 limit where $k_{close} << k_{chem}$, every opening event produces a concerted exchange for all exposed amide protons with the rate equal to the opening rate, $k_{obs} = k_{open}$. Unfortunately, we cannot use the $k_{close} >> k_{chem}$ criterion to test for EX2 behavior as commonly employed in site-resolved NMR studies [(1)] because we lack explicit information on $k_{close}$ for BtuB. However, we can take advantage of the behavior of HDX/MS isotopic envelopes to identify whether our measurements are in the EX1 or EX2 limit. Specifically, in the EX1 limit, the concerted exchange should produce a decreasing protonated isotopic envelope and a concomitant rise in the corresponding deuterated isotopic envelope [(3)]. In the EX2 limit, however, the isotopic envelope for a peptide should exhibit a continuous increase in mass as a function of time, as exchange at each residue in the peptide is uncorrelated. The D uptake is consistent with EX2 behavior for the presented peptides, and we therefore interpret the data as occurring in the EX2 limit.

**Supplemental Material and Methods**

**Bacterial strains and plasmids.**

TonB 32-239 in pHis Parallel and full length BtuB in pAG1 were received as a generous gift from Professor Robert Nakamoto at the University of Virginia. pHis is derived from pET22b and contains a 0.7kb insertion of TonB except for the amino terminal transmembrane helix, described further in [(4)]. pAG1 is derived from pUC8 and contains a 2.4kb insertion of the BtuB gene from *E. coli* [(5)]. For construction of the LH4 plasmid, QuikChange II mutagenesis was used to insert a five additional histidines, a glycine, and two serines to form a hexahistidine tag and GSS spacer between residues Asp448 and Thr450 using the following primers: (Position 449 is already a histidine in the wild-type sequence)

forward 5'-CATCATCATCATCACGGCAGCAGCACCCTGAAATATTACAACGAAGGG-3' and

reverse 5'- CCGTGATGATGATGATGGTGATCATCATAATCGATCAAATCAC-3'.

**Protein expression and purification.**

$TonB_{\Delta N}$ (residues 32-239) containing an amino terminal HisTag and proTEV cleavage site was transformed into BL21 (DE3) *E. coli* on LB plates supplemented with 100 μM ampicillin

(LB/AMP). A single colony was picked to inoculate an overnight culture of 10 mL of 2 x YT medium. This culture was used to inoculate 1L of 2 x YT medium and grown to 0.7-0.8 OD at 37 ºC before induction with 0.5 mM IPTG at 20 ºC for 5 hours. Cells were harvested by centrifugation at 6500 RPM for 12 minutes at 4 ºC, and frozen or resuspended in 25 mM Tris-HCl, 100 mM NaCl, 1 mM PMSF, pH 7.5 and lysed immediately by 5 passes through a French press. The lysate was clarified by centrifugation for 30 min at 12,5000 RPM at 4 ºC in a Sorvall Legend X1R centrifuge fitted with a FiberLite F15-8x50c and applied to a 5 mL column of equilibrated nickel-NTA resin. The column was washed with 20 column volumes (100 mL) of wash buffer: 50 mM Tris-HCl, 100 mM NaCl, 20 mM imidazole, pH 7.5 and eluted 6 times with 5 mL each of elution buffer, which was the same as wash buffer, except that the imidazole concentration was increased to 300 mM. Protein containing fractions were pooled and dialyzed in a length of 3.5 kDa molecular weight cutoff tubing against 2 L of dialysis buffer: 50 mM Tris-HCl, 100 mM NaCl, pH 7.6 for 2 hours before addition of 1 mg of homemade TEV protease in 1 mL of TEV protease storage buffer: 50 mM Tris-HCl, 1 mM EDTA, 5 mM DTT, 50% glycerol, 0.01% Trion, pH 7.6. TEV cleavage was allowed to proceed under dialysis for 16 hours at 4 ºC. Cleavage was assessed by SDS-PAGE electrophoresis and a subsequent round of subtractive Ni-IMAC performed to remove uncleaved protein. The flowthrough from this was concentrated and injected onto a 5 mL pre-packed HiTrap SP HP anion exchange column (Cytiva, Washington D.C., Cat# 17115201) for purification by a 20 min gradient from 100% buffer A (25 mM tris, 50 mM NaCl, pH 7.5) to 100% buffer B (25 mM tris, 1 M NaCl, pH 7.5). Protein-containing fractions were checked by SDS-PAGE and concentrated to 30-70 µM in 25mM tris 50 mM NaCl pH 7.5 using spin filtration before addition to pelleted outer membrane stocks to obtain BtuB:TonB$_{ΔN}$ or BtuB:B12:TonB$_{ΔN}$ complex samples (with B12 addition as above) with a final TonB concentration of 26 µM.

BtuB wild-type or LH in pAG1 was transformed into either BL21 *E. coli* or a specialized strain with four abundant OMPs deleted, termed ΔABCF [(6)]. A single colony was picked to inoculate 5mL of LB/AMP and grown overnight at 30 ºC with shaking at 225 RPM. The next day, 1 mL of this culture was taken to inoculate 50 mL of LB/AMP medium and grown at 30 ºC for 6 – 8 hours, before seeding 1 L of cell culture. This 1 L of culture was either grown to $OD_{600}$ = 0.6 – 0.8 and induced or split into up to 10 L of LB/AMP medium for large-scale expression. Protein was induced at 30 ºC for 4 hours with 1 mM IPTG. Cells were harvested by centrifugation in a Sorvall RC6+ centrifuge equipped with F10s 6x500y rotor for 30 minutes at 6000 RPM (6340 x g) at 4 ºC and frozen at -80 ºC until purification.

Frozen cells were thawed on ice and resuspended in 40 mL of resuspension buffer per liter of cell culture. The resuspension buffer composition was: 50 mM Tris, 100 mM NaCl, pH 8.0, supplemented before lysis with 10 mM $MgCl_2$, 0.1 mg/mL lysozyme (Sigma, ref# 6876), 0.05 mg/mL DNAse I (Goldbio, St. Louis, MO, cat# D-300-1). For cell pellets grown in ΔABCF *E. coli*, $MgCl_2$ was omitted from the resuspension buffer as it causes clumping of cells and outer membranes. Resuspended cells were kept on ice and homogenized by at least 5 passes through an Emulsiflex-C5 French press set to 40 psi. The cell lysate was clarified by centrifugation in a Sorvall Legend-X1R centrifuge equipped with F15-8x50cy rotor for 30 minutes at 7000 RPM at 4 ºC. The clarified lysate was ultracentrifuged in 70 mL polycarbonate (Beckman Coulter, Brea, CA, part # 355655) bottles for 60 minutes at 40,000 RPM (~100,000 x g) at 4 ºC in a Beckman Coulter Optima L-100 XP ultracentrifuge equipped with 45 Ti rotor. The resulting total membrane pellets were washed by resuspension with a number 10 round tipped paint brush in ~6 mL per gram of cell membrane mass of resuspension buffer and Dounce homogenization before ultracentrifugation, performed as before. Washed total membranes were resuspended and Dounce homogenized as before prior to addition of 0.5% sodium N-lauroyl sarcosinate (“sarkosyl”, from a 10% w/v stock in water) and incubation on a rotisserie for 30 minutes to 1 hour at room temperature or overnight at 4 ºC to solubilize the inner membranes. Following another ultracentrifugation step as described above, outer membranes were again homogenized and ultracentrifuged as described above to wash out residual sarkosyl. This step was repeated once more prior to final resuspension of the washed outer membrane pellet in 5 mL of resuspension buffer. Resuspended washed outer membranes were aliquoted and stored at 4 $^{o}$C for use in native-like outer membrane hydrogen exchange experiments, or frozen at -20 $^{o}$C until extraction for use in detergent hydrogen exchange experiments.

Extraction from outer membranes was performed by addition of concentrated n-octyl tetraoxyethylene ($C_8E_4$, BACHEM, Bubendorf, Switzerland, article # 4006356.0025) to a final concentration of 150 mM (4.6% w/v, or 19 x CMC) and incubation on a rotisserie at room temperature for 2 hours. Insoluble debris was removed by ultracentrifugation for 60 minutes at 100,000 x g (47,246 RPM) at 4 $^{o}$C in a Beckman Coulter MAX-XP microultracentrifuge equipped with TLA-55 rotor. $C_8E_4$-extracted BtuB was either further purified by IMAC on a 5 mL Ni-NTA column (Goldbio, cat# H-320-100), with a 50-600 mM LiCl gradient on a DEAE FastFlow anion exchange column with 20mM BisTris pH 6.9 or taken directly for detergent exchange. Detergent exchange was carried out by incubation of BtuB aliquots with 50 x CMC of lauryl maltose neopentyl glycol (LMNG, Anatrace, Maumee, OH, ref# NG310, added from a concentrated stock in water) for 1 hour at room temperature before injection on a BioRad Enrich 650 SEC 10 x 300 mm column with 20 mM BisTris pH 6.9 as the running buffer. BtuB-containing fractions were supplemented with 1.5 x CMC of dodecyl maltoside (DDM, Anatrace, D322S, added from concentrated stock in water) immediately upon elution and either concentrated or immediately taken for HXMS experiments.

**Hydrogen Exchange Labeling Experiments.**

Hydrogen exchange label, simultaneous quench/digestion, and injection steps were all performed manually for experiments performed on protein in native-like outer membranes. Labeling in a solution of ~ 93%D at room temperature (22 $^{o}$C) was initiated by addition of 2 µL of thoroughly agitated native-like outer membrane stock (in 20 mM BisTris pH 6.9) to 28 µL of deuteration buffer (50 mM NaPi, 150 mM NaCl, made with dibasic sodium hydrogen phosphate and adjusted to $pD_{read}$ = $pD_{desired}$ – 0.4 using DCl). For labeling reactions involving B12 liganded samples, 1% v/v of 100x concentrated stocks (2 mM B12 and 100 mM $CaCl_2$) were added to aliquots both protein stock and label solutions to obtain a final concentration of 20 µM B12 and 1 mM $CaCl_2$. For labeling reactions involving TonB CTD, aliquots of BtuB samples were allowed to settle and half of the volume was aspirated and replaced with a solution of $TonB_{\Delta N}$ concentrated to 50 µM in 20 mM BisTris pH 6.9. Immediately after addition of outer membrane suspensions to label buffer, samples were triturated 5 or more times to ensure complete mixing. Quenching of the HX label reaction and digestion was performed by repeated trituration of the HX label reaction with a 200 µL pipette and aspiration of the entire volume and transfer into an Eppendorf tube containing 37 µL freshly prepared quench mix sitting inside an ice-chilled heat transfer block: 30 µL of 600 mM glycine, 4 M urea, pH 2.5, 2.0 µL of thrice-desalted 10 mg/mL porcine pepsin (Sigma-Aldrich, ref# P6887) in 100 mM sodium citrate, pH 4.4, 2.5 µL of 2.5 mM DDM in water, and 2.5 µL of a 300 mg/mL aqueous suspension of $ZrO_2$ coated silica (beads extracted from Supelco brand cartridges, Sigma-Aldrich, ref# 55261-U). Digestion was carried out for 3 minutes before resuspension by repeated trituration and aspiration of the whole ~67 µL volume onto a cellulose acetate spin cup (Thermo Pierce, Waltham, MA ref# 69702) held in a 1.5 mL polypropylene Eppendorf tube. The quenched digestion mix was filtered by spinning for 30 seconds at 16,000 RCF in an Eppendorf 5415 R centrifuge fitted with a FA-45-24-11 rotor pre-chilled to 4 $^{o}$C. Filtered peptide-containing solutions were aspirated with a glass ice-chilled 50 µL glass Hamilton syringe and immediately injected into the LC sample loading loop.

For peptide assignment MS/MS runs, the protocol above was repeated, except for a brief incubation in an undeuterated “mock HX” buffer using the same samples as the HX experiments and with 8M urea solubilized BtuB derived from inclusion bodies instead of native membranes. For these, the “mock HX” and quench buffers’ urea concentrations were varied from 0 to 4M to optimize peptide coverage and signal intensity. Additionally, in-exchange controls to account for deuteration towards the 41.5%D level present in the quench solution were performed by mixing HX and quench buffers prior to addition of BtuB. Similarly, full-exchange (“All D”) controls were performed by incubating 8M-urea solubilized unfolded BtuB with $pD_{corr}$ 7.2 label buffer for 600 – 64500 seconds at room temperature. In-exchange controls measured deuteration levels between 0 – 5%D, and All D controls measured deuteration levels of 70-75%D. The mean level of back exchange in peptides used in our analysis was 27%, with 73% of the label remaining on the longest urea control. There was a weak correlation of back exchange levels with sequence position, with peptides covering residues 1-79 having an average back exchange level of 25 and peptides covering residues 80-137 having an average back exchange level of 30%.

Fluid handling operations for HXMS experiments in detergents were carried out with a Trajan LEAP HDX PAL workstation. 25 to 30 μM of DDM-solubilized BtuB wild-type or “LH4” samples (“LH4” is a construct of BtuB with a hexahistidine tag inserted after position 449) were diluted into >99%D 25 mM NaPi, 25 mM NaOAc, 100 mM NaCl, pH 5.0 – 7.0 labeling buffer for 0.25 - 900 minutes at 22 $^{o}$C before 2-fold dilution into a 4 $^{o}$C-chilled vial containing HX quench buffer: 600 mM glycine pH 2.5, 2 M urea, with the quench vials being manually supplemented with 2 μL of 10 mg/mL pepsin at the start of each time course. The quench reaction time was set to 1 minute, and each fluid transfer operation involving BtuB was followed with two cycles of aspiration and dispensing to ensure proper mixing.

**Mass Spectrometry.**

For labeling experiments, upon injection peptides were trapped and desalted by flowing across a 5x1 mm C8 5 μm particle column (TARGA brand, Higgins Analytical, Mountain View, CA, TP-M501-C085) for 3 minutes at 100 μL/min. After desalting, the trap was diverted to be in line with a Dionex Ultimate-3000 gradient pump flowing at 20 μL/min, with a 50x0.5 μm C18 3 μm particle column (TARGA brand, Higgins Analytical, TS-05M5-C183) in line for peak refocusing upstream of the HESI-II (Thermo Fisher Scientific, cat# IQLAAEGABBFACNMAGY) probe attached to a Thermo Q Exactive mass spectrometer, and a 14-minute gradient from 10% to 60% buffer B was immediately started. MS data were collected from 2 to 13 minutes into the gradient, and the flow diverted to waste otherwise. Buffer A was 0.1% formic acid in water, and buffer B was 0.1% formic acid in acetonitrile. After the main gradient, the pump was set to 90% B for 2 minutes, then down to 10% B for a minute, and two trapezoid washes up to 90% B were performed to clean the columns before the next injection. Electrospray ionization was performed at 180 $^{o}$C, with the following parameters: spray voltage set to 3.2 kV, 1 microscan per scan, resolution 140,000, AGC target 3e6, minimum IT 100 ms, scan range 400-2000 m/z, dynamic exclusion 10 ppm, sheath gas flow rate 5, aux and sweep gas flow rates 0.

For assignment experiments, peptides were analyzed by performing unlabeled injections of outer membrane samples, or an 8 M urea solubilized concentrated stock of BtuB purified from inclusion bodies, after being expressed without a signal sequence. MS/MS data were acquired with the following parameters: data collected from 2 to 13 minutes of an identical gradient to HX runs detailed above, Full MS as detailed above, dd-$MS^2$: resolution: 17,500, automatic gain correction target: 1e5, maximum integration time: 200 ms, loop count: 10, MSX count: 1, TopN: 10, isolation window: 1.9, m/z isolation offset: 0.00, scan range: 200 to 2000 m/z, fixed first mass: 100.0 m/z, NCE / stepped: NCE 27, spectrum data type: profile.

**Data Processing.**

Peptide signals were assigned by searching against a sequence database containing pepsin and BtuB sequences using the meta-search program SearchGUI version 3.3.21, (search settings: unspecific cleavage, precursor charge 1-8, isotopes 0-1, precursor m/z tolerance 10.0 ppm, fragment m/z tolerance 0.5 Da, no post-translational modifications, peptide length 5-30) incorporating X! Tandem, MS Amanda, and Comet algorithms. Data were inspected in PeptideShaker version 1.16.45, and results were imported into HDExaminer 3.1 (Sierra Analytics, Modesto, CA). HDExaminer was used to fit isotopic envelopes, enabling bimodal fits where separately exchanging subpopulations were present. Integration bounds of retention time as well as m/z were adjusted manually after an initial round of calculations performed by the program. While attempting to maintain consistency, isotopic envelope integration bounds were sometimes overridden to exclude obvious noise and carried over peptide signal near 0%D. Occasionally, integration bounds had to be manually chosen to achieve consistent extraction of heavily overlapped envelopes in cases where a peptide's independently exchanging subpopulations failed to diverge at early time points.

A subsequent round of manual adjustment was performed considering inter-peptide differences, to ensure that all overlapping peptides for a sequence fragment exhibited the same behavior. In cases where a peptide grossly disagreed with others covering the same sequence it was excluded from further analysis. In some instances, all time points from pairs of peptides with similar m/z signals and LC elution times were manually re-integrated multiple times to assess the contributions from overlapping envelopes.

**Proteolysis Assay.**

BtuB used for proteolysis assays was either washed and resuspended unextracted outer membranes, $C_8E_4$ extraction reaction supernatants, or Ni-IMAC and/or SEC purified DDM-solubilized protein fractions. BtuB was diluted to 1 µM on ice in HX quench buffer: 600 mM glycine pH 2.5 buffer, with 0 - 3 M urea, and the reactions were started by adding 0.5 mg/mL pepsin from a concentrated stock. At times ranging from 7 seconds to 243 minutes aliquots of the reaction were diluted with an equal volume of proteolysis quench buffer: 2 M Tris pH 9.8 and spun at 15,000 x g for 3 minutes before addition of Laemmli sample buffer and boiling prior to loading on SDS-PAGE. Electrophoresis was carried out in Novex Wedgewell 4-20% acrylamide gradient Tris-Glycine mini gels (Thermo Fisher Scientific, Waltham, MA), run in an Invitrogen Mini Gel tank at 180 V for 80 minutes at 4 $^{o}$C with ice packed around the cell. Gels were stained with AcquaStain protein stain (Bulldog Bio), Coomassie R-250, Invitrogen His-Tag In-Gel Stain, or transferred to western blots, and either imaged or selected bands were cut out for downstream applications such as Edman degradation or trypsin/endopeptidase LysC digestion followed by LC/MS/MS identification of proteolysis products.

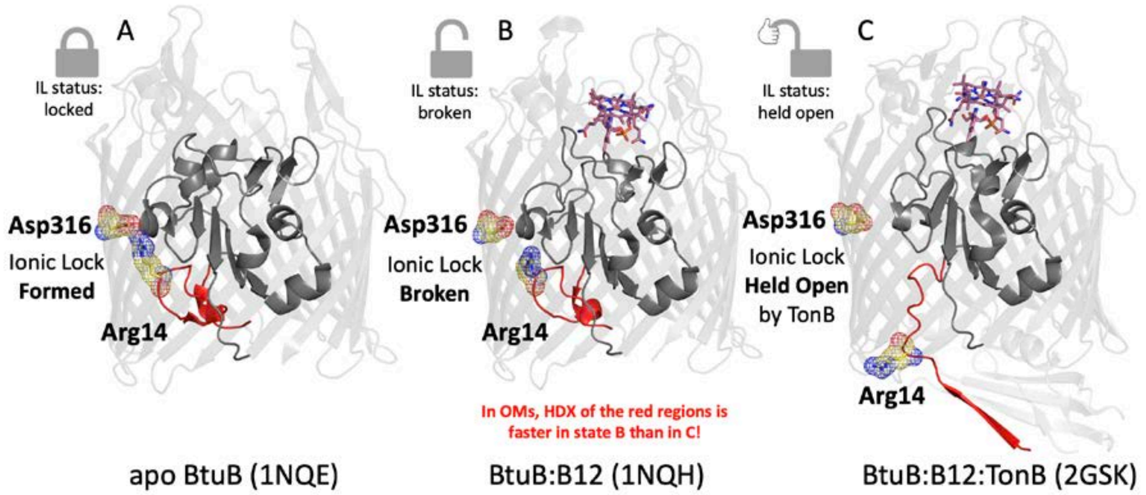


**Fig. S1. Detailed comparison of IL and ILR in PDB structures. A)** apo BtuB (1NQE) **B)** BtuB:B12 complex (1NQH) C) BtuB:B12:TonB$_{CTD}$ complex (2GSK) structures. The amino terminal tail through the ILR is colored red. The IL is shown as yellow sticks and mesh, and B12 is shown as pink sticks, with non-carbon atoms colored by element.

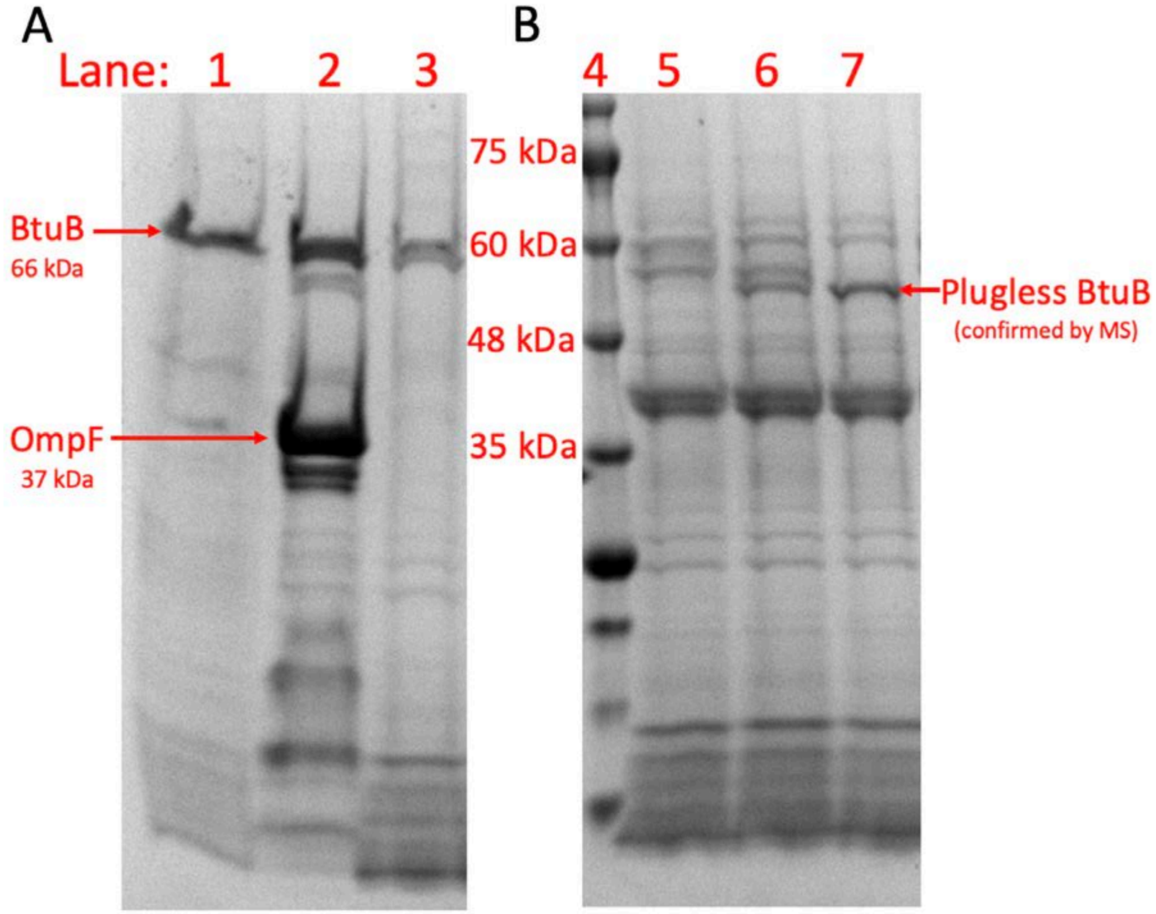


**Fig. S2. SDS-PAGE of A) OMP abundance and B) soluble pepsin digestion of BtuB in OMs. A)** Total OMP content of the three OM samples used in this study. Lanes 1 and 3, duplicates prepared from OMs of **Δ**ABCF E. coli, that lack several naturally abundant OMPs, e.g., OmpF. Lane 2 is prepared from OM of wild-type E. coli. **B)** Digestion of BtuB in OM by pepsin under HDX quench conditions, with lanes 5, 6, and 7 corresponding to digestion times of 1, 3, and 10 minutes, respectively. Lane 4 is a molecular weight marker. Note the progressive weakening of the intact BtuB band and the appearance of a plugless BtuB band (whose identity was confirmed by tryptic digest experiments performed at an independent facility), along with several intermediates. A digestion time of 3 minutes was used for HDX samples.

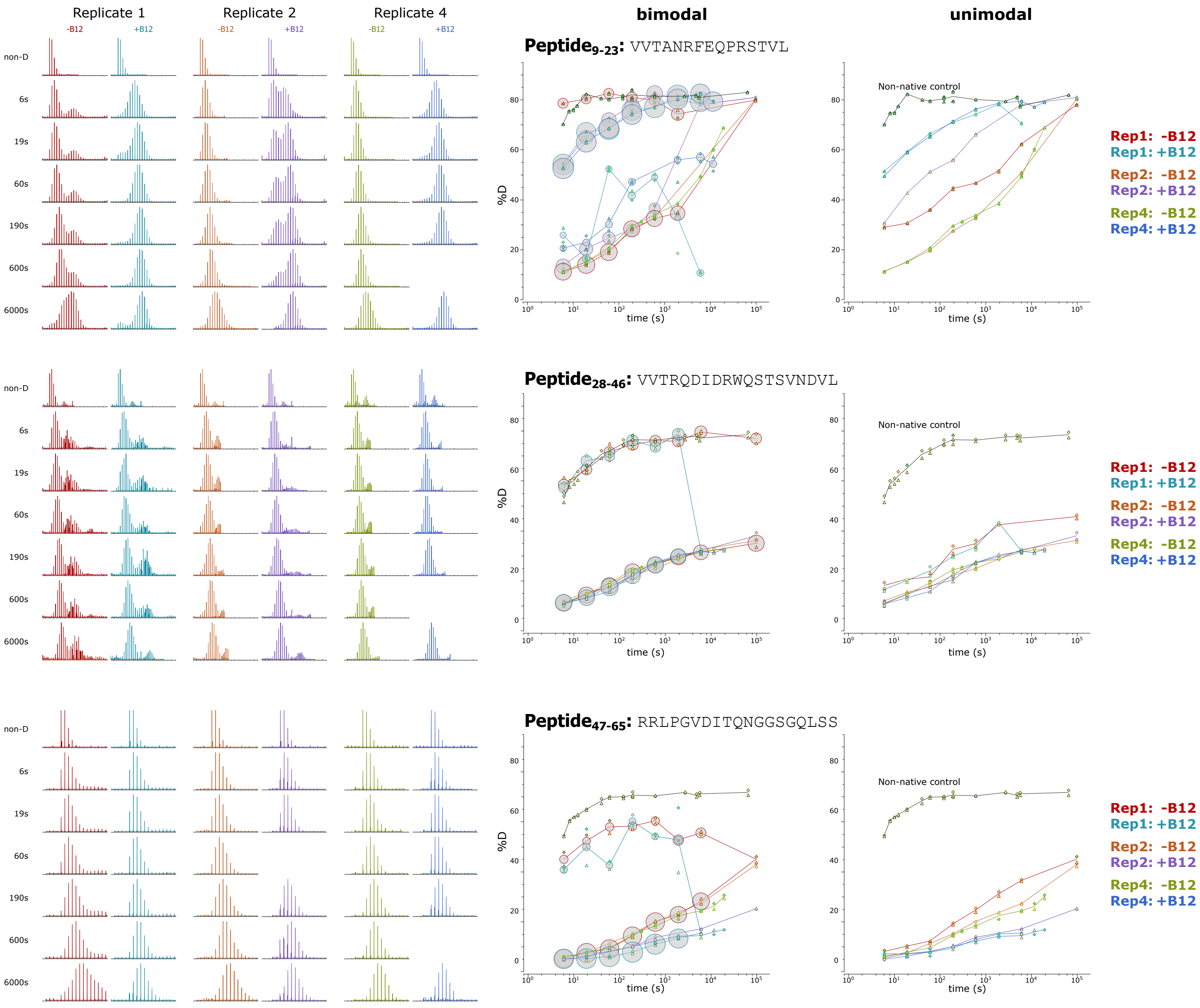


**Fig. S3. Bimodal exchange indicates sample heterogeneity.** Many peptides' mass spectra exhibit bimodal behavior, consistent within (but not across) bio-replicates. The raw data (left column) contains distinctly exchanging isotopic envelopes, which occasionally match non-native control data (black). Utilizing bimodal analysis (middle column) produces a more accurate estimation of the acceleration in deuterium uptake upon B12 binding compared to the unimodal analysis (right column).

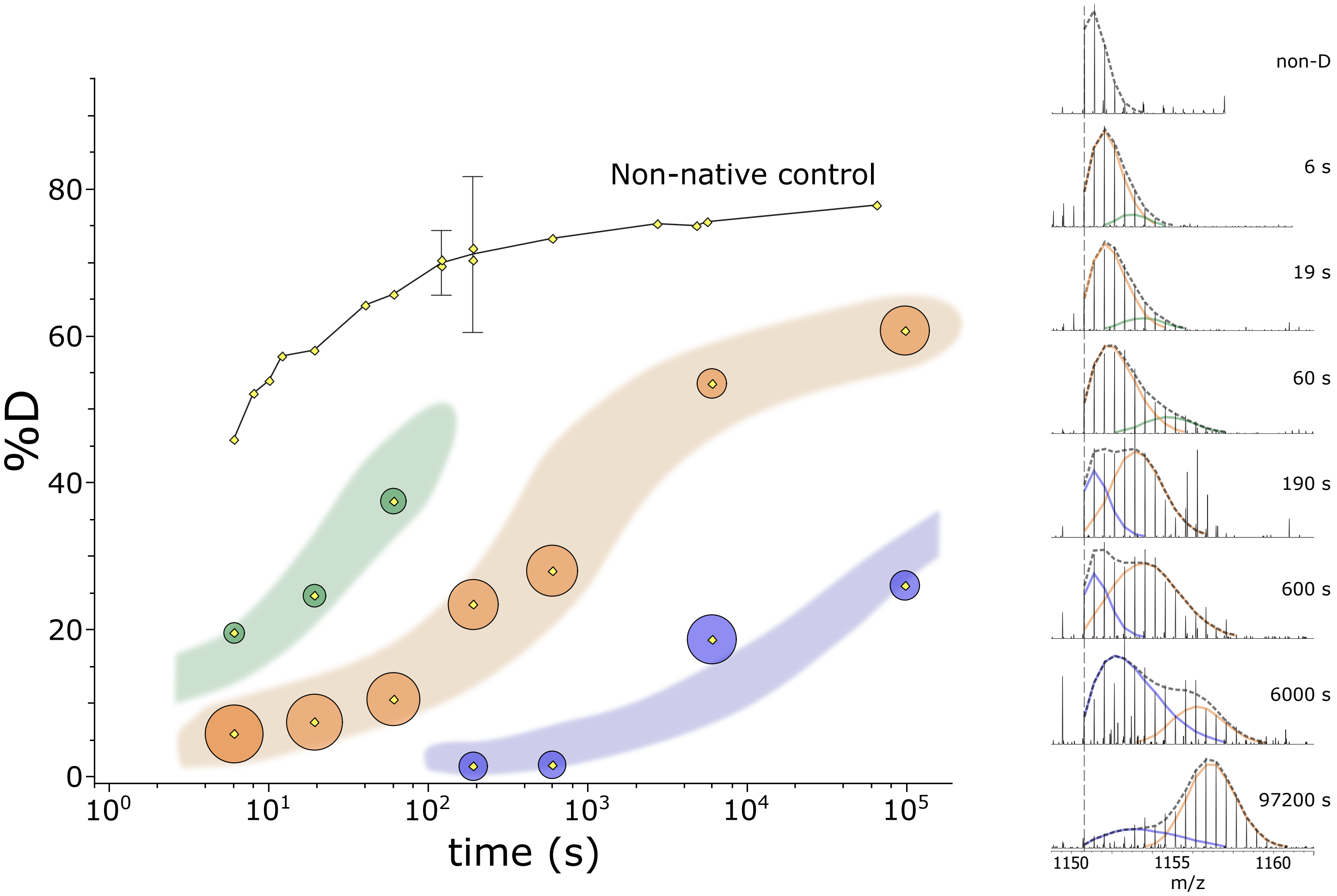


**Fig. S4. Multi-modal uptake in SB3 loop. Left:** The data for bioreplicate 2 of Peptide$_{80\text{-}102}$ contains at least three distinctly exchanging species, labeled by color, and are plotted along with data from a non-native control (black line). **Right:** Extracted peptide m/z spectra with traced bimodal fits, following the same coloring scheme. From 60s to 190s, a switch occurs regarding which pair of species is being identified with the bimodal analysis. At a given time point, HDExaminer approximates the mass spectrum with (at most) a bimodal fit, so one subpopulation is lost. For example, the slowest-exchanging (blue) species is not fit and therefore merged with the middle (orange) distribution from fits for labeling times < 100 sec and the same is true for the fastest (green species) for fits of data sampled at labeling times > 100 sec.

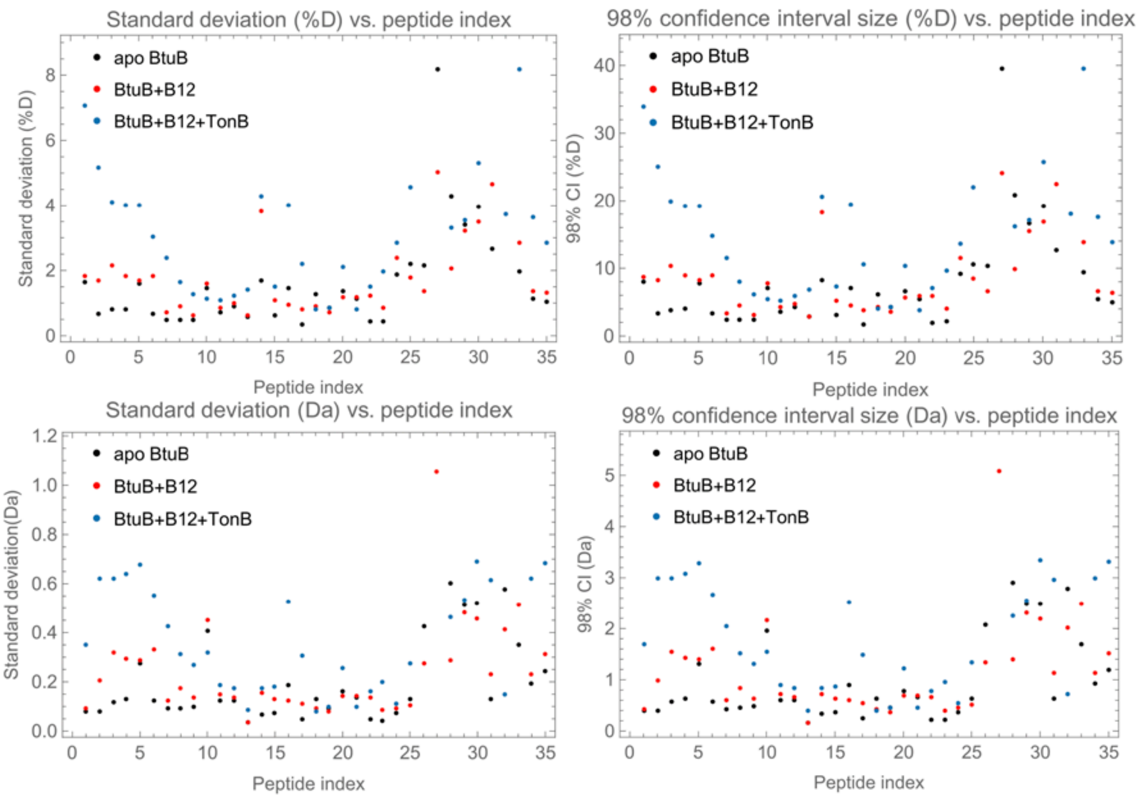


**Fig. S5. Statistical summary plots.** Scatterplots show the reproducibility as standard deviations (left) or 98% confidence intervals (right), in terms of %D (top) or Daltons (bottom). Peptides are ordered sequentially by amino terminus. Peptide index 26, where reproducibility begins to decrease, corresponds to $Peptide_{80\text{-}102}$, the first to cover the SB3 loop apex.

Fig S6

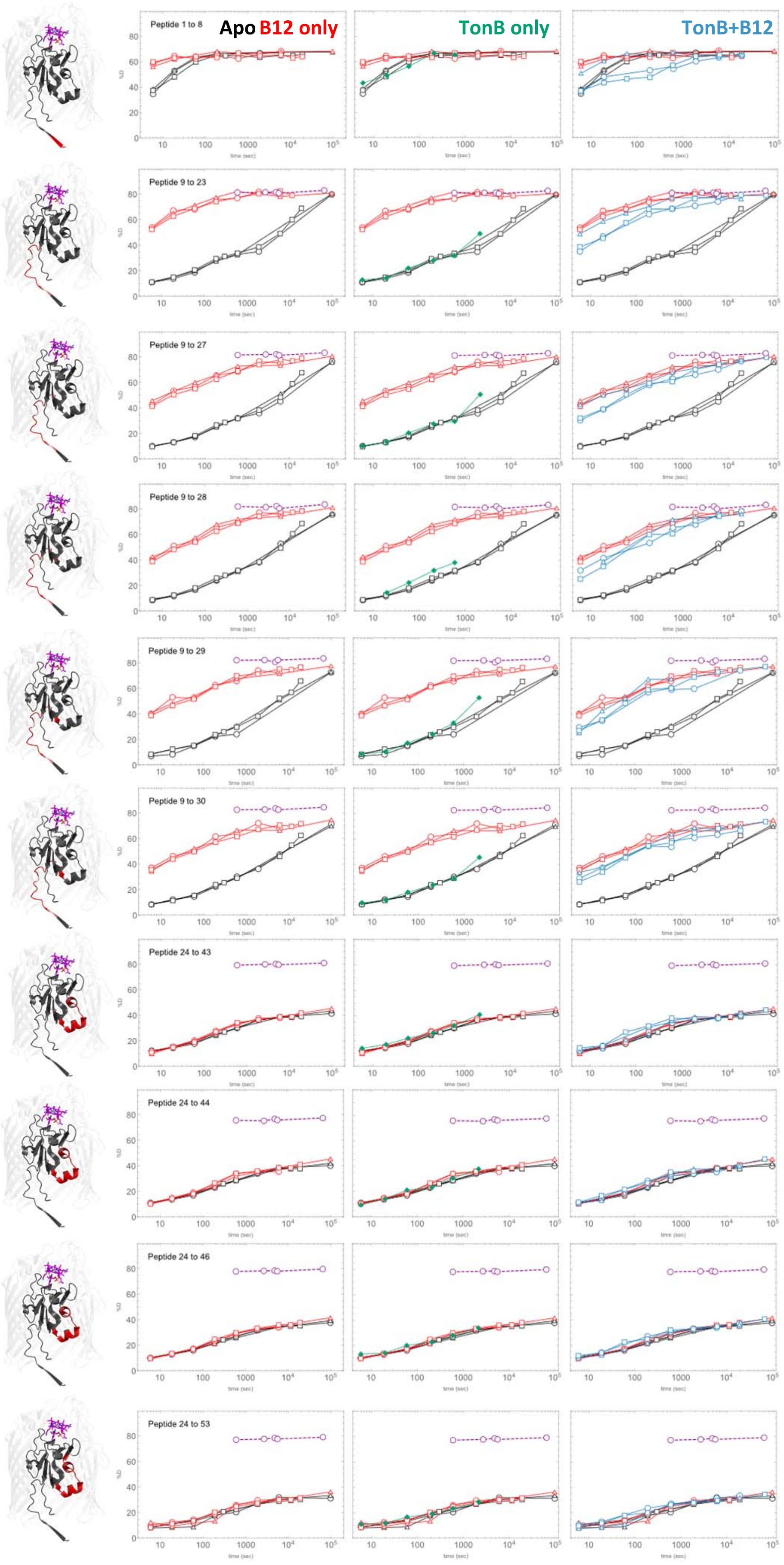
Apo B12 only
TonB only
TonB+B12
Peptide 1 to 8
Peptide 9 to 23
Peptide 9 to 27
Peptide 9 to 28
Peptide 9 to 29
Peptide 9 to 30
Peptide 24 to 43
Peptide 24 to 44
Peptide 24 to 46
Peptide 24 to 53

Fig S6 (cont)

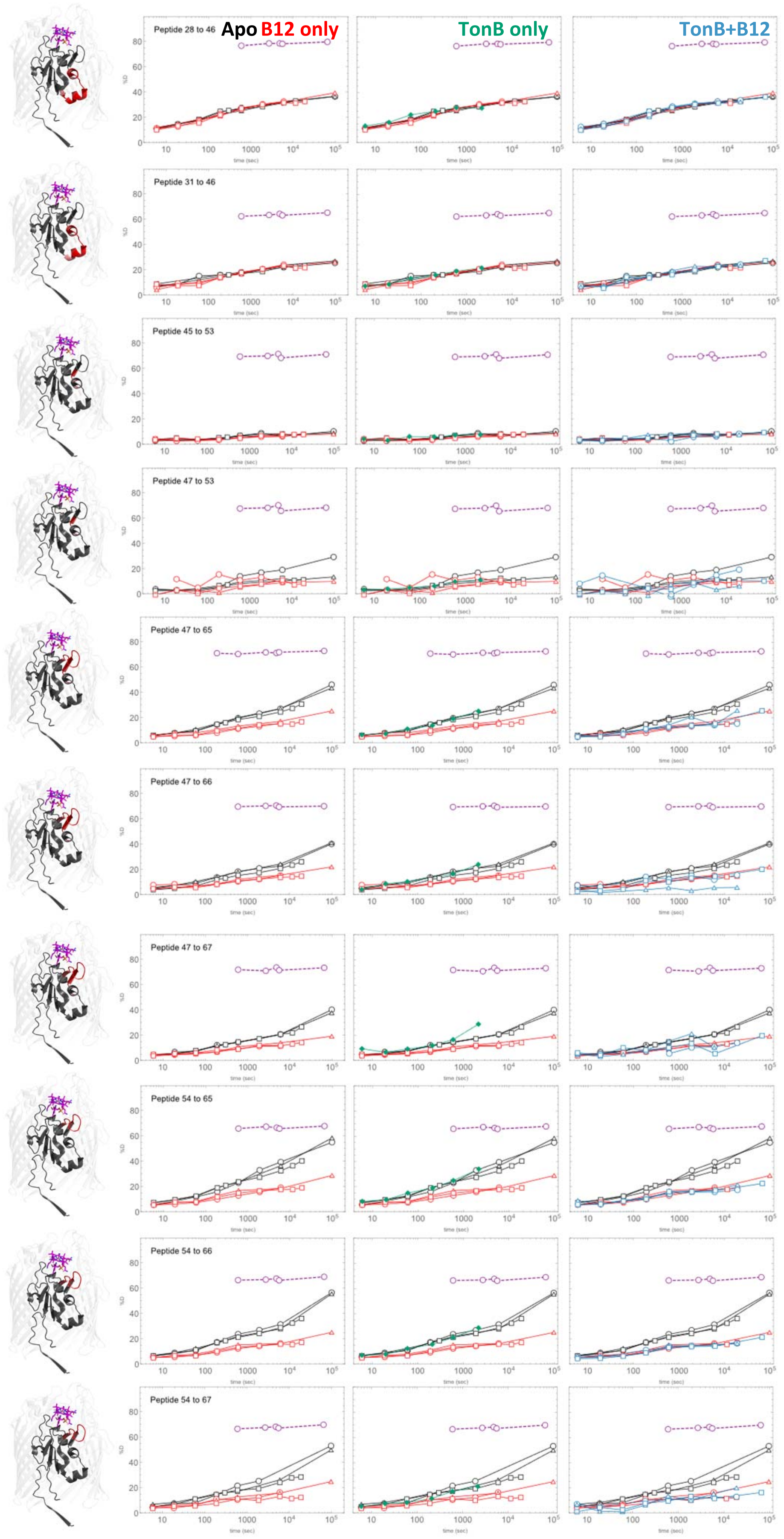
Apo
B12 only
TonB only
TonB+B12
Peptide 28 to 46
Peptide 31 to 46
Peptide 45 to 53
Peptide 47 to 53
Peptide 47 to 65
Peptide 47 to 66
Peptide 47 to 67
Peptide 54 to 65
Peptide 54 to 66
Peptide 54 to 67

Fig S6
(cont)

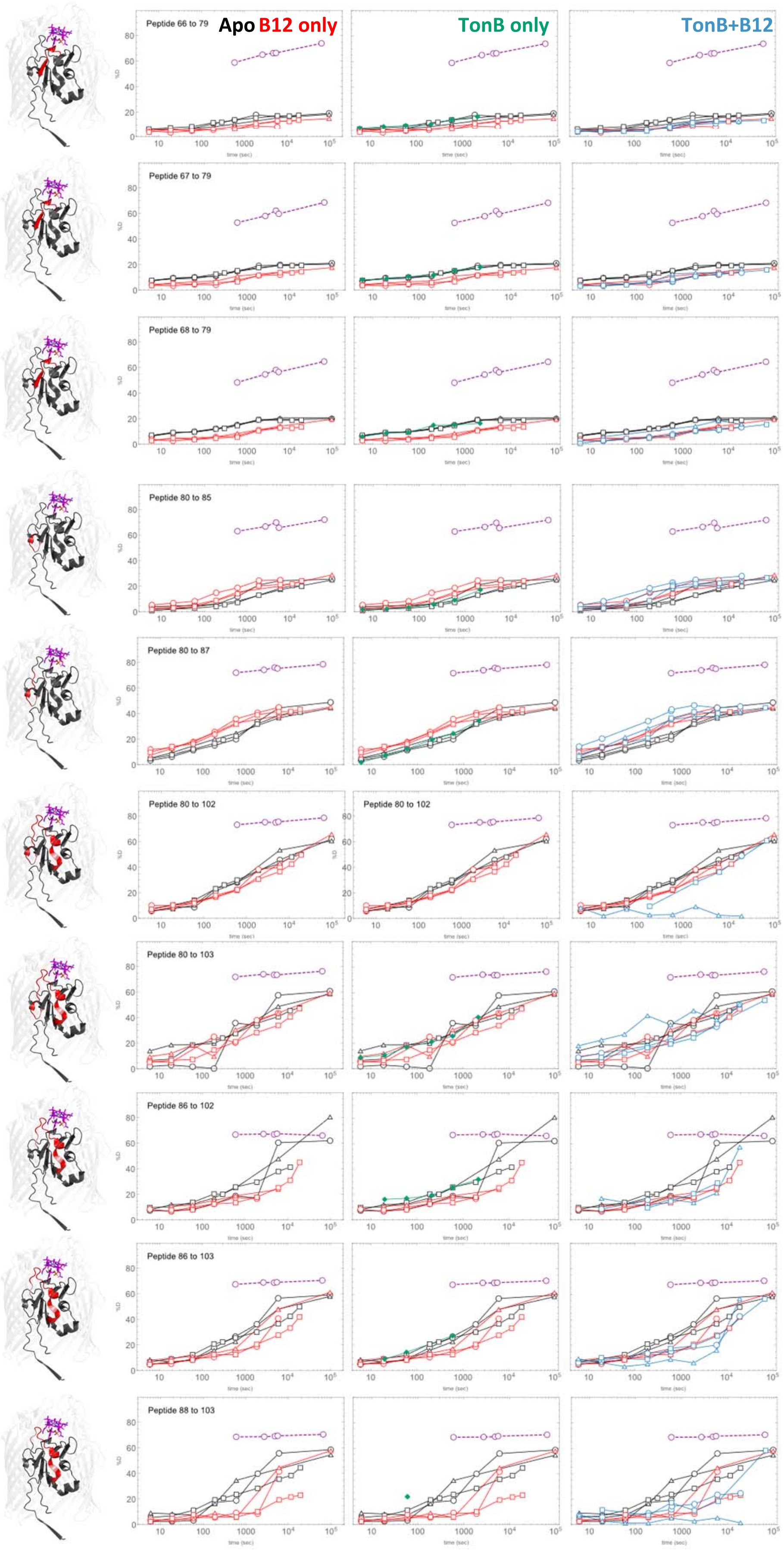

Apo B12 only
TonB only
TonB+B12
Peptide 66 to 79
Peptide 67 to 79
Peptide 68 to 79
Peptide 80 to 85
Peptide 80 to 87
Peptide 80 to 102
Peptide 80 to 103
Peptide 86 to 102
Peptide 86 to 103
Peptide 88 to 103

Fig S6 (cont)

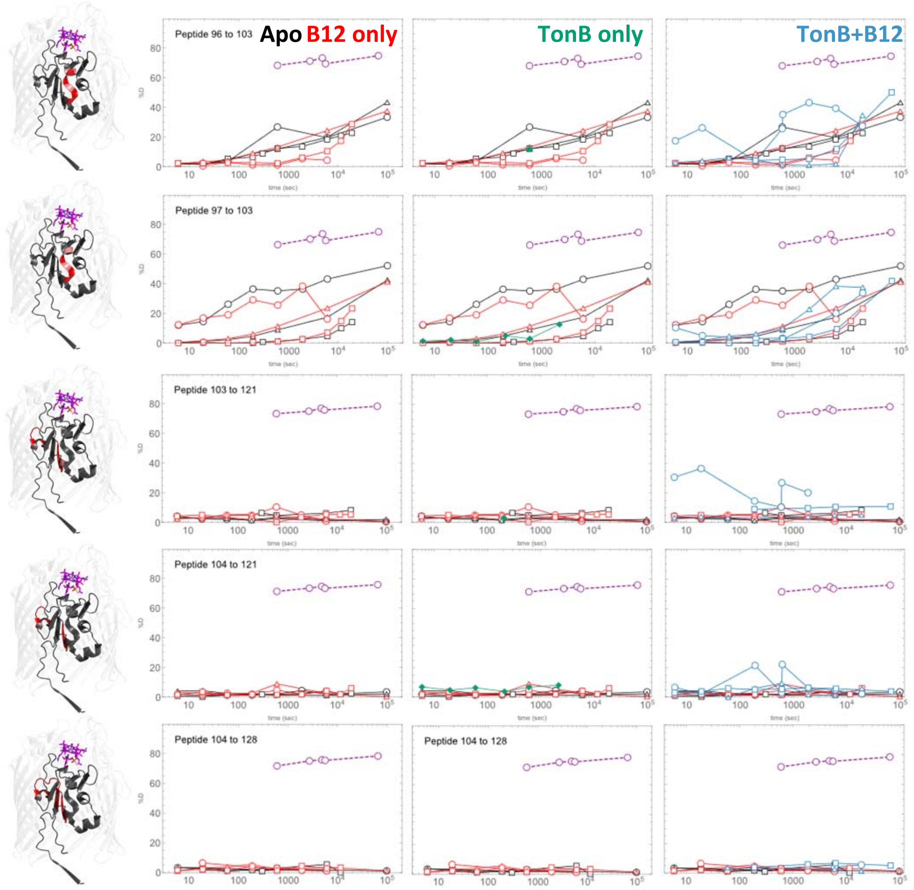


**Fig. S6.1 through S6.35. Deuterium uptake plots.** Three deuterium uptake plots are shown for each peptide, together with a structure highlighting the peptide in red. The first two residues of a peptide, which are non-observable due to rapid back exchange, are colored a brighter shade of red to distinguish them. Deuterium uptake plot color scheme is as follows: Apo BtuB (black), BtuB+B12 (red), BtuB+B12+TonB (blue), BtuB+TonB, (green), and nonnative BtuB only control (purple). For clarity, only the last 5 points for the nonnative control are shown.

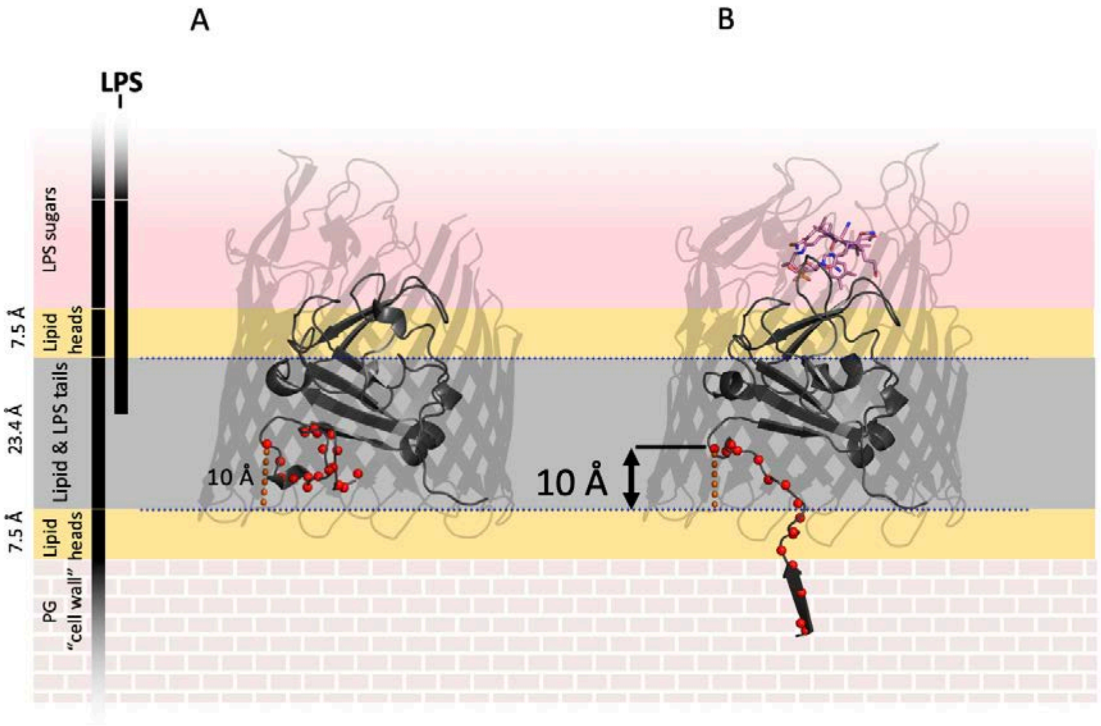


**Fig. S7. Outer membrane environment.** Structures of **A)** apo BtuB (PDBID: 1NQE) and **B)** BtuB:B12:TonB ternary complex (PDBID: 2GSK) highlighting amide nitrogen atoms of residues 6-23 as red spheres. The 10 Å distance from Leu23 to the membrane plane defined by OPM [7] is shown as a dashed orange line. The shaded background shows approximate zones of various outer membrane layers. Dark grey indicates lipid tails, yellow indicates the headgroup region, solid pale red indicates extracellular lipopolysaccharide, and brick indicates peptidoglycan (the cell wall). The lipid zone thickness value of 23.4 Å is taken from OPM, the headgroup value of 7.5 Å is taken from [8] although the model presented here with parallel planes is an oversimplification. The membrane plane has been measured to be significantly distorted around BtuB in lipid vesicles as the hydrophobic thickness varies significantly around the circumference of the β-barrel. However, regardless of membrane plane tilt, several amides on the amino terminus exit the lumen into the periplasmic zone occupied by peptidoglycan where $[OD^-]$ apparently is lower than in bulk solvent, based on our findings that HDX for this region remains $10^2$ slower than $k_{chem}$ in the presence of B12.

**Table S1.** HDX peptide statistical summary table. Standard deviations, number of triplicated points in each condition, and sizes of 98% confidence interval sizes are provided for each peptide. Some peptides in conditions with TonB present were excluded due to poor signal quality and are marked with NAN.

| Peptide Index | Peptide Length | Start | End | Number of observable sites | Standard deviation (S.D.) of apo points (%D) | Number of triplicated apo points | S.D. of BtuB+B12 points (%D) | Number of triplicated BtuB+B12 points | S.D. of BtuB+B12+TonB points (%D) | Number of triplicated BtuB+B12+TonB points | S.D. of BtuB+B12+TonB points (%D) | S.D. of apo points (Da) | S.D. of BtuB+B12 points (Da) | S.D. of BtuB+B12+TonB points (Da) | %D in last urea point (64500 sec) (Taken as 100 - %back exchange) | 98% Conf. interval size for apo state (Da) | 98% C.I. size for BtuB+B12 state (Da) | 98% C.I. size for BtuB+B12+TonB state (Da) |
|---|---|---|---|---|---|---|---|---|---|---|---|---|---|---|---|---|---|---|
| 1 | 8 | 1 | 8 | 5 | 1.65 | 6 | 1.81 | 6 | 7.05 | 5 | 7.05 | 0.08 | 0.09 | 0.35 | NAN | 0.40 | 0.44 | 1.70 |
| 2 | 15 | 9 | 23 | 12 | 0.69 | 5 | 1.69 | 5 | 5.16 | 5 | 5.16 | 0.08 | 0.20 | 0.62 | 83.03 | 0.40 | 0.98 | 2.99 |
| 3 | 19 | 9 | 27 | 15 | 0.80 | 5 | 2.14 | 5 | 4.12 | 5 | 4.12 | 0.12 | 0.32 | 0.62 | 83.32 | 0.58 | 1.55 | 2.99 |
| 4 | 20 | 9 | 28 | 16 | 0.83 | 6 | 1.85 | 6 | 3.99 | 4 | 3.99 | 0.13 | 0.30 | 0.64 | 83.63 | 0.64 | 1.43 | 3.09 |
| 5 | 21 | 9 | 29 | 17 | 1.61 | 5 | 1.69 | 6 | 4.00 | 4 | 4.00 | 0.27 | 0.29 | 0.68 | 83.58 | 1.32 | 1.39 | 3.28 |
| 6 | 22 | 9 | 30 | 18 | 0.67 | 6 | 1.84 | 6 | 3.06 | 5 | 3.06 | 0.12 | 0.33 | 0.55 | 84.54 | 0.58 | 1.60 | 2.66 |
| 7 | 20 | 24 | 43 | 18 | 0.50 | 6 | 0.69 | 6 | 2.37 | 5 | 2.37 | 0.09 | 0.13 | 0.43 | 80.99 | 0.43 | 0.60 | 2.06 |
| 8 | 21 | 24 | 44 | 19 | 0.49 | 6 | 0.91 | 6 | 1.64 | 5 | 1.64 | 0.09 | 0.17 | 0.31 | 77.49 | 0.45 | 0.84 | 1.51 |
| 9 | 23 | 24 | 46 | 21 | 0.47 | 6 | 0.64 | 6 | 1.28 | 5 | 1.28 | 0.10 | 0.13 | 0.27 | 79.72 | 0.48 | 0.64 | 1.30 |
| 10 | 30 | 24 | 53 | 28 | 1.45 | 6 | 1.61 | 6 | 1.14 | 4 | 1.14 | 0.41 | 0.45 | 0.32 | 79.13 | 1.96 | 2.17 | 1.54 |
| 11 | 19 | 28 | 46 | 17 | 0.72 | 6 | 0.86 | 6 | 1.10 | 5 | 1.10 | 0.12 | 0.15 | 0.19 | 79.66 | 0.59 | 0.71 | 0.90 |
| 12 | 16 | 31 | 46 | 14 | 0.88 | 4 | 0.98 | 6 | 1.23 | 4 | 1.23 | 0.12 | 0.14 | 0.17 | 65.19 | 0.60 | 0.66 | 0.83 |
| 13 | 9 | 45 | 53 | 6 | 0.60 | 6 | 0.60 | 6 | 1.42 | 5 | 1.42 | 0.04 | 0.04 | 0.09 | 71.20 | 0.17 | 0.17 | 0.41 |
| 14 | 7 | 47 | 53 | 4 | 1.69 | 6 | 3.81 | 5 | 4.28 | 5 | 4.28 | 0.07 | 0.15 | 0.17 | 68.38 | 0.33 | 0.74 | 0.83 |
| 15 | 19 | 47 | 65 | 12 | 0.63 | 6 | 1.07 | 6 | 1.51 | 4 | 1.51 | 0.08 | 0.13 | 0.18 | 72.81 | 0.36 | 0.62 | 0.87 |
| 16 | 20 | 47 | 66 | 13 | 1.44 | 6 | 0.95 | 6 | 4.03 | 5 | 4.03 | 0.19 | 0.12 | 0.52 | 70.10 | 0.91 | 0.59 | 2.53 |
| 17 | 21 | 47 | 67 | 14 | 0.37 | 4 | 0.80 | 6 | 2.18 | 5 | 2.18 | 0.05 | 0.11 | 0.31 | 73.37 | 0.25 | 0.54 | 1.48 |
| 18 | 12 | 54 | 65 | 10 | 1.29 | 6 | 0.90 | 6 | 0.82 | 5 | 0.82 | 0.13 | 0.09 | 0.08 | 67.80 | 0.62 | 0.44 | 0.40 |
| 19 | 13 | 54 | 66 | 11 | 0.86 | 6 | 0.72 | 6 | 0.87 | 5 | 0.87 | 0.10 | 0.08 | 0.10 | 69.01 | 0.46 | 0.38 | 0.46 |
| 20 | 14 | 54 | 67 | 12 | 1.35 | 5 | 1.17 | 6 | 2.13 | 5 | 2.13 | 0.16 | 0.14 | 0.26 | 69.76 | 0.78 | 0.68 | 1.23 |
| 21 | 14 | 66 | 79 | 12 | 1.12 | 6 | 1.20 | 6 | 0.81 | 5 | 0.81 | 0.13 | 0.14 | 0.10 | 74.01 | 0.65 | 0.70 | 0.47 |
| 22 | 13 | 67 | 79 | 11 | 0.42 | 6 | 1.24 | 6 | 1.49 | 4 | 1.49 | 0.05 | 0.14 | 0.16 | 68.70 | 0.22 | 0.66 | 0.79 |
| 23 | 12 | 68 | 79 | 10 | 0.44 | 6 | 0.85 | 6 | 1.99 | 2 | 1.99 | 0.04 | 0.09 | 0.20 | 64.84 | 0.21 | 0.41 | 0.96 |
| 24 | 6 | 80 | 85 | 4 | 1.89 | 6 | 2.38 | 6 | 2.84 | 5 | 2.84 | 0.08 | 0.10 | 0.11 | 72.18 | 0.37 | 0.46 | 0.55 |
| 25 | 8 | 80 | 87 | 6 | 2.20 | 6 | 1.77 | 6 | 4.57 | 5 | 4.57 | 0.13 | 0.11 | 0.27 | 78.64 | 0.64 | 0.51 | 1.33 |
| 26 | 23 | 80 | 102 | 20 | 2.14 | 6 | 1.39 | 6 | 21.19 | NAN | NAN | 0.43 | 0.28 | NAN | 78.74 | 2.07 | 1.34 | NAN |
| 27 | 24 | 80 | 103 | 21 | 8.18 | 6 | 5.01 | 5 | 10.17 | 4 | 10.17 | 1.72 | 1.05 | 2.14 | 76.42 | 8.30 | 5.08 | 10.32 |
| 28 | 17 | 86 | 102 | 14 | 4.50 | 6 | 2.06 | 4 | 3.53 | NAN | NAN | 0.63 | 0.29 | NAN | 66.09 | 3.04 | 1.40 | NAN |
| 29 | 18 | 86 | 103 | 15 | 3.42 | 6 | 3.21 | 6 | 3.53 | 3 | 3.53 | 0.51 | 0.48 | 0.53 | 70.58 | 2.48 | 2.32 | 2.56 |
| 30 | 16 | 88 | 103 | 13 | 3.98 | 5 | 3.51 | 6 | 5.33 | 4 | 5.33 | 0.52 | 0.46 | 0.69 | 70.63 | 2.50 | 2.20 | 3.34 |
| 31 | 8 | 96 | 103 | 5 | 4.27 | 5 | 4.66 | 5 | 12.30 | 3 | 12.30 | 0.21 | 0.23 | 0.62 | 74.92 | 1.03 | 1.13 | 2.97 |
| 32 | 7 | 97 | 103 | 4 | 14.39 | 6 | 10.38 | 6 | 3.74 | 4 | 3.74 | 0.58 | 0.42 | 0.15 | 75.04 | 2.78 | 2.00 | 0.72 |
| 33 | 19 | 103 | 121 | 18 | 1.95 | 2 | 2.87 | 3 | 8.16 | NAN | NAN | 0.35 | 0.52 | NAN | 78.38 | 1.69 | 2.49 | NAN |
| 34 | 18 | 104 | 121 | 17 | 1.14 | 6 | 1.37 | 5 | 3.64 | 5 | 3.64 | 0.19 | 0.23 | 0.62 | 75.84 | 0.93 | 1.13 | 2.98 |
| 35 | 25 | 104 | 128 | 24 | 1.02 | 4 | 1.37 | 4 | 2.86 | NAN | NAN | 0.25 | 0.33 | NAN | 78.40 | 1.19 | 1.58 | NAN |
| Average of peps 1-35 | 16.49 | | | 13.60 | 2.00 | 5.54 | 2.00 | 5.63 | 3.99 | 4.48 | 3.35 | 0.24 | 0.24 | 0.40 | 75.08 | 1.15 | 1.16 | 1.94 |
| Average of peps 1-25 | 16.08 | | | 13.00 | 1.00 | 5.68 | 1.37 | 5.88 | 2.60 | 4.64 | 2.60 | 0.12 | 0.17 | 0.31 | 75.33 | 0.58 | 0.80 | 1.49 |

**Dataset 1.** HDX data table **(separate .xlsx file).** HDX data and all computed quantities used the analysis.
